# Identification of Dysregulated Pathways and key genes in Human Retinal Angiogenesis using Microarray Metadata

**DOI:** 10.1101/2020.08.30.273870

**Authors:** Umadevi Subramanian, Bharanidharan Devarajan

## Abstract

Retinal angiogenesis is a common neovascularization mechanism that causes severe irreversible vision loss in the number of retinal diseases worldwide. Patients often do not respond to the current anti-angiogenic therapies and have a vision loss. Understanding the various angiogenic pathways and factors involved in the pathogenic mechanism is vital for disease management. In this study, to identify dysregulated angiogenic pathways and specific angiogenic factors involved in vision-threatening diseases namely proliferative diabetic retinopathy (PDR), retinopathy of prematurity (ROP) and neovascular age-related macular degeneration (nAMD), we downloaded microarray metadata of samples and obtained the differentially expressed genes (DEGs) in all the disease and each disease samples compared to controls. Subsequently, we performed Gene Set Enrichment (GESA) analysis for pathways, a protein-protein interaction (PPI), and angiome network analysis using R and Cytoscape software. We identified highly enriched dysregulated pathways that were neuroactive ligand receptor interaction and cytokine-cytokine receptor interaction. The angiogenic–associated DEGs were predominately related to the cytokine-cytokine receptor interaction pathway, which we further confirmed with RNA-seq data of PDR samples. Together, our analysis of these data elucidated the molecular mechanisms of retinal angiogenesis and provided potential angiogenic targets for therapeutics.

## 1. Introduction

Retinal angiogenesis is the common dysregulated pathway of numerous blinding disorders, namely, proliferative diabetic retinopathy (PDR), retinal artery (or vein) occlusion, retinopathy of prematurity (ROP), sickle cell retinopathy^1^, and neovascular age-related macular degeneration (nAMD)^2^. Although there are subtle differences in the mechanism of angiogenesis, they are characterized as common neovascularization event. Angiogenesis is a physiological process through which new blood vessels formed from pre-existing vessels under hypoxic conditions^3^.Hypoxia-inducible-1α drives the expression of substantial pro-angiogenic growth factors and induces endothelial cells into a tip cell phenotype. The tip cell proliferates slowly and activates matrix metalloproteinases (MMPs), plasmin, uPA, and tPA, and cleaving numerous extracellular matrix (ECM) proteins. The activated MMPs and other proteins stimulate the Notch ligand in tip cells and proliferate further to elongate a new vessel^4,5^. As the newly formed vessels mainly serve a role in a wound healing response, they usually do not restore tissue integrity, but instead, cause visual impairment^6,7^.

Vascular endothelial growth factor (VEGF) is the major growth factor, induced in hypoxic conditions. VEGF family members VEGF-A, VEGF-B, VEGF-C, VEGF-D and placental growth factor (PIGF), and their receptors are the validated regulators of angiogenic signaling pathway^8^. VEGF-A is the potent growth factor and mediator of intraocular neovascularization. VEGF-A stimulates not only the development of new blood vessels, but also, microaneurysm formation, capillary occlusion with ischemia, and promoting increased vascular permeability^9^. VEGF-B can regulate the function of endothelial cells. VEGF-C and VEGF-D increase vascular permeability as well as induce angiogenesis. PIGF plays a role in pathogenic angiogenesis by increasing the activity of VEGF-A^10^.

The current anti-angiogenic agents, approved by the Food and Drug Administration, inhibit the VEGF pathway^8^. The anti-VEGF drugs are the molecules that target VEGF isoforms, or inhibiting VEGF receptors, or inhibiting VEGF downstream signaling^11^. About 30% of PDR patients fail to respond to initial treatment, and the majority of the responders would require multiple rounds of intravitreal injections^12^. Therefore, anti-VEGF treatment is potentially required for years, and chronic VEGF inhibition may cause side effects and drug resistance. The potential consequence of VEGF blockade is hypertension, proteinuria, impairment of wound healing/collateral vessel development, and inhibition of bone growth, infertility, inhibition of skeletal muscle, regeneration, cardiac remodeling and neuroprotective effect in the ischaemic retina^13^. Also, a dose-dependent decrease in ganglion cells was reported in rats^14^.

Studies show that VEGF independent factors may stimulate angiogenesis, particularly in tumors^15^. Zhang et al. suggested identifying the angiogenic factors other than VEGF for a better therapeutic approach^16^. Gene expression studies have been widely performed to identify angiogenic factors. However, such studies will provide numerous key genes that are differentially expressed, and thus, it is a challenging task to detect reliable targets. On the molecular level, a combination of the differential gene expression (which may vary in each individual) is assumed to dysregulate the common cellular pathway^17^. Therefore, identification of the dysregulated pathways is essential for understanding disease mechanisms and the identification of targets for future application of custom therapeutic decisions^18^. The accumulation of large amounts of gene expression data in public databases provides an opportunity for identifying commonly dysregulated pathways and the gene targets that are differentially expressed^19^.

In the present study, we aim to identify the dysregulated angiogenic pathways in addition to VEGF dependent angiogenic pathway in retinal diseases. To achieve our aim, we designed this study with several steps that are (i) to combine the microarray data from different reliable experiments and adjusting batch effects, (ii) to identify robust differentially expressed genes (DEGs), (iii) to conduct the pathway enrichment analysis, (iv) to identify key dysregulated angiogenic pathways and angiogenic factors through network analysis and (v) to compare and confirm the identified dysregulated angiogenic pathways and angiogenic factors with available RNA-Seq data

## 2. Materials and Methods

### 2.1. Gene expression data and preprocessing

All the microarray samples used in this study were systematically searched and downloaded from NCBI-GEO^20^ after the manual curation of the sample details. Probe IDs were mapped to official gene symbols in the DAVID database^21^, and non-protein coding genes, as defined by an assignment of HUGO gene symbol, were excluded from the study. In case of genes with multiple probes, gene expression level was aggregated from probeset to gene-level data by taking the maximum value across all the samples^22^.

### 2.2. Missing value imputation and normalization of data

All the samples selected for the study were combined by their gene symbol using R script. The missing gene expression values were imputed by Bayesian Principal Component Analysis (BPCA) method^23^, which performed well even with low entropy. Further, the whole data were quantile normalized^24^ to make the distribution of values in all the samples into the same. The outlier samples were removed based on the principal component analysis (PCA) plot. Further it was confirmed with Euclidean distance-based clustering using R script.

### 2.3. Identification of differentially expressed genes (DEGs) and pathway enrichment

DEGs were selected from the list by filtering, only if the moderated t-test FDR was ≤ 0.05 and the fold change (FC - a difference of the means between two groups when all the values were log2 transformed) ≥ 2 (up-regulated) or ≤ −2 (down-regulated). For the pathway analysis, Gene Set Enrichment Analysis (GSEA) proposed by Subramanian et al. implemented with GenePattern tools was performed^25^. The DEGs and FC values were regarded as a pre-ranked list, and c2.cp.kegg.v6.1 was interrogated as pathway database. We used default parameters and set the minimum gene set size 2. The output enrichment score reflects the degree to which a gene set in the pathway is over-represented at the extreme (low or high) of the entire ranked list of FC. For the pathway enrichment, normalized enrichment score (NES) was considered as a metric of influence of DEGs with p-value ≤ 0.05.

Analysis was carried out with different perspectives, differential gene expression and pathway regulation between retinal angiogenesis (case) samples and controls, case and tissue (retina, and RPE/C) specific controls, cases and disease (dry AMD, non-Proliferative Diabetic Retinopathy - NPDR, and Diabetic Mellitus - DM) specific control samples, disease specificity (nAMD, PDR, ROP) and control samples as given in figure 1. Among the DEGs, the angiogenesis-related genes were identified by GO term in GSEA and the list is given by Chu et al.^26^. Further, the STRING database was used to retrieve the predicted interactions for the identified DEGs and graphically represented using Cytoscape 3.2.0.

**Figure 1.**
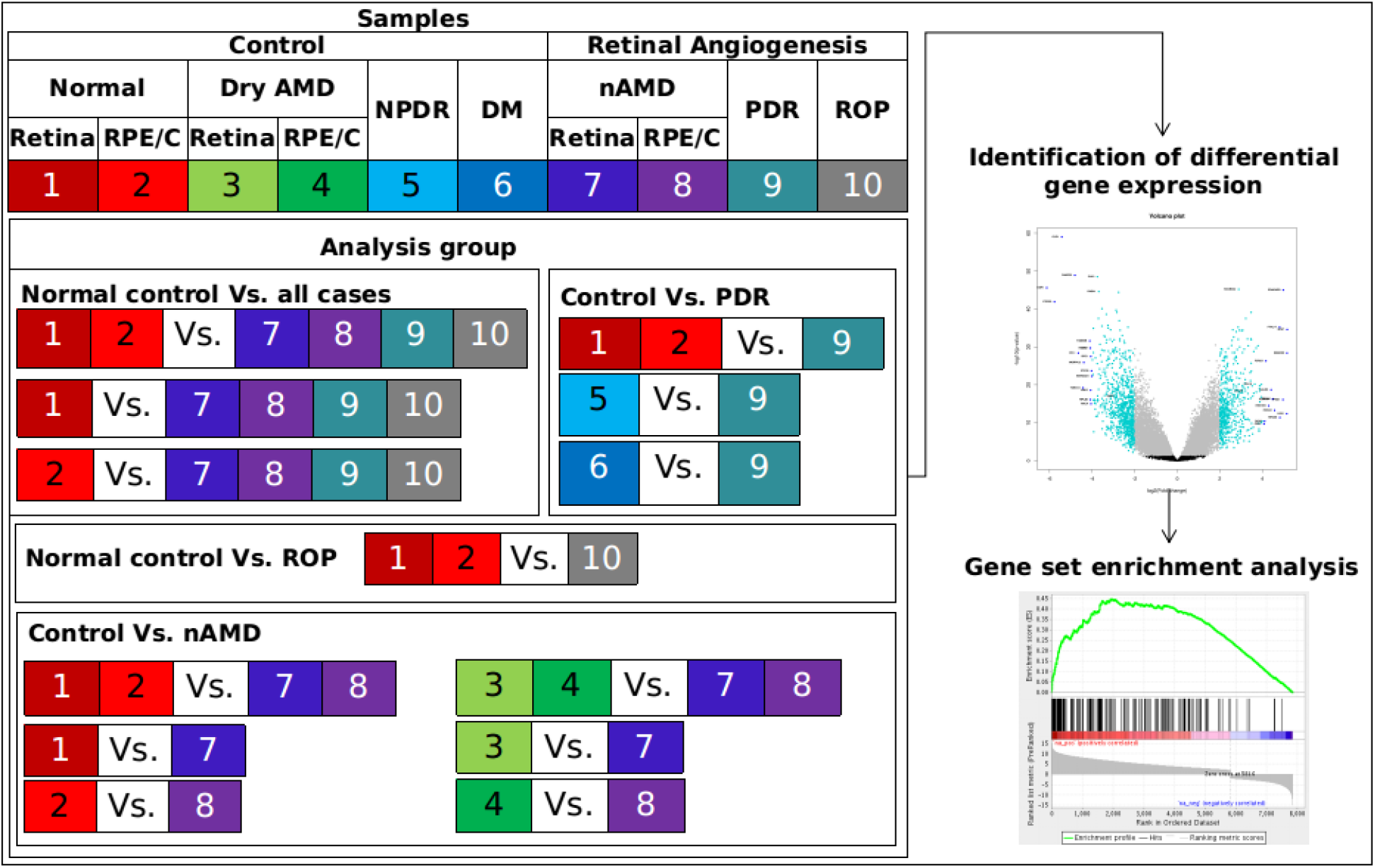
Schema of the workflow - samples, analysis groups used, and the methods followed

### 2.4. RNA-Seq analysis

The available RNA-Seq samples related to this study in NCBI-SRA are PDR (Accession No.: SRP097696) and whole retinal (SRP119766) control samples. They were downloaded by SRA Toolkit, aligned to human reference genome hg38 using CLC Genomics Workbench 12.0, quantile normalized in R, and analyzed using EdgeR.

## 3. Results

### 3.1. Standardization of gene expression data

The analysis was primarily done with 243 control samples, 95 normal retinal control (GSE60436 (3 samples), GSE28133 (19), GSE29801 (55), GSE32614 (12), GSE41019 (3), GSE57864 (3)), 138 normal RPE/C control (GSE18811 (31), GSE29801 (96), GSE43257 (2), GSE50195 (7), GSE5741 (2)), and 32 disease cases includes, 16 (8 retina, and 8 RPE/C) nAMD (Accession No.: GSE29801), 6 fibrovascular membrane of PDR (GSE60436), and 10 retinal microvascular endothelial cells of ROP (GSE5946) samples. The studies with disease specific control samples include, 64 dry AMD retina (GSE29801 (61), GSE17938 (3)), 9 dry AMD RPE/C (GSE50195), 6 neural retinal NPDR (GSE53257) and 6 neural retinal DM (GSE53257) samples. After preprocessing and merging of data, 15.75% of missing data were imputed. All the samples which contain 18621 genes were quantile normalized (Figure S1). The 318 control samples (95 retina, 138 RPE/C) used in the primary study were obtained from different technical platforms. Eventhough they were quantile normalized, to reduce the bias the outlier samples have to be removed. In order to this, all the disease samples and control samples were subjected to PCA (Figure S2). The disease samples were retained, the control samples which were scatterd/ outgrouped were removed, and clustering tree was construted with remaining samples. This process was iteratively done once observed the clear clustering of case and control samples (Figure S3). This results to 31 retina and 34 RPE/C control samples.

### 3.2. Dysregulated pathways of all the cases with all normal control samples

One thousand three hundred eighty-three genes displayed differential expression with two-fold change (FDR ≤ 0.05) in all the cases compared to normal control samples (Figure S4). Of these, 852 were downregulated and 531 upregulated. Gene set enrichment analysis (GSEA) with respect to pathways showed that 237 DEGs were associated with 22 different dysregulated pathways (significant with p ≤ 0.05, NES of pathways are given in Table 1). We then analyzed DEGs that are associated with angiogenesis. We identified 127 angiogenic-associated enriched DEGs, of which 31 were based on gene ontology (GO) (NES: 1.74 with p = 0.01) and 96 DEGs were based on the literature. The highly enriched pathways with their significantly altered DEGs and their association with angiogenesis were graphically represented in Figure 2.

**Table 1.** List of dysregulated pathways enriched in retinal angiogenic disease samples compared to normal control samples.

| Dysregulated pathway | Normalized |  |
| --- | --- | --- |
|  | enrichment score | p value |
| Neuroactive ligand receptor interaction | 3.101 | 0.000 |
| Cytokine cytokine receptor interaction | 2.553 | 0.000 |
| Arachidonic acid metabolism | 1.931 | 0.007 |
| Aminoacyl tRNA biosynthesis | -1.572 | 0.042 |
| Tight junction | -1.631 | 0.042 |
| Pathogenic Escherichia coli infection | -1.646 | 0.032 |
| Cysteine and methionine metabolism | -1.667 | 0.035 |
| Vibrio cholerae infection | -1.753 | 0.018 |
| Endocytosis | -1.768 | 0.014 |
| Regulation of actin cytoskeleton | -1.872 | 0.012 |
| Epithelial cell signaling in helicobacter pylori infection | -1.873 | 0.016 |
| Arrhythmogenic right ventricular cardiomyopathy | -1.886 | 0.011 |
| p53 signaling pathway | -1.892 | 0.008 |
| Protein export | -1.909 | 0.004 |
| Lysosome | -1.925 | 0.004 |
| Adherens junction | -2.040 | 0.002 |
| Spliceosome | -2.094 | 0.004 |
| Alzheimers disease | -2.108 | 0.000 |
| Parkinsons disease | -2.324 | 0.002 |
| Huntingtons disease | -2.494 | 0.000 |
| Oxidative phosphorylation | -2.579 | 0.000 |
| Ribosome | -4.047 | 0.000 |

**Figure 2.**
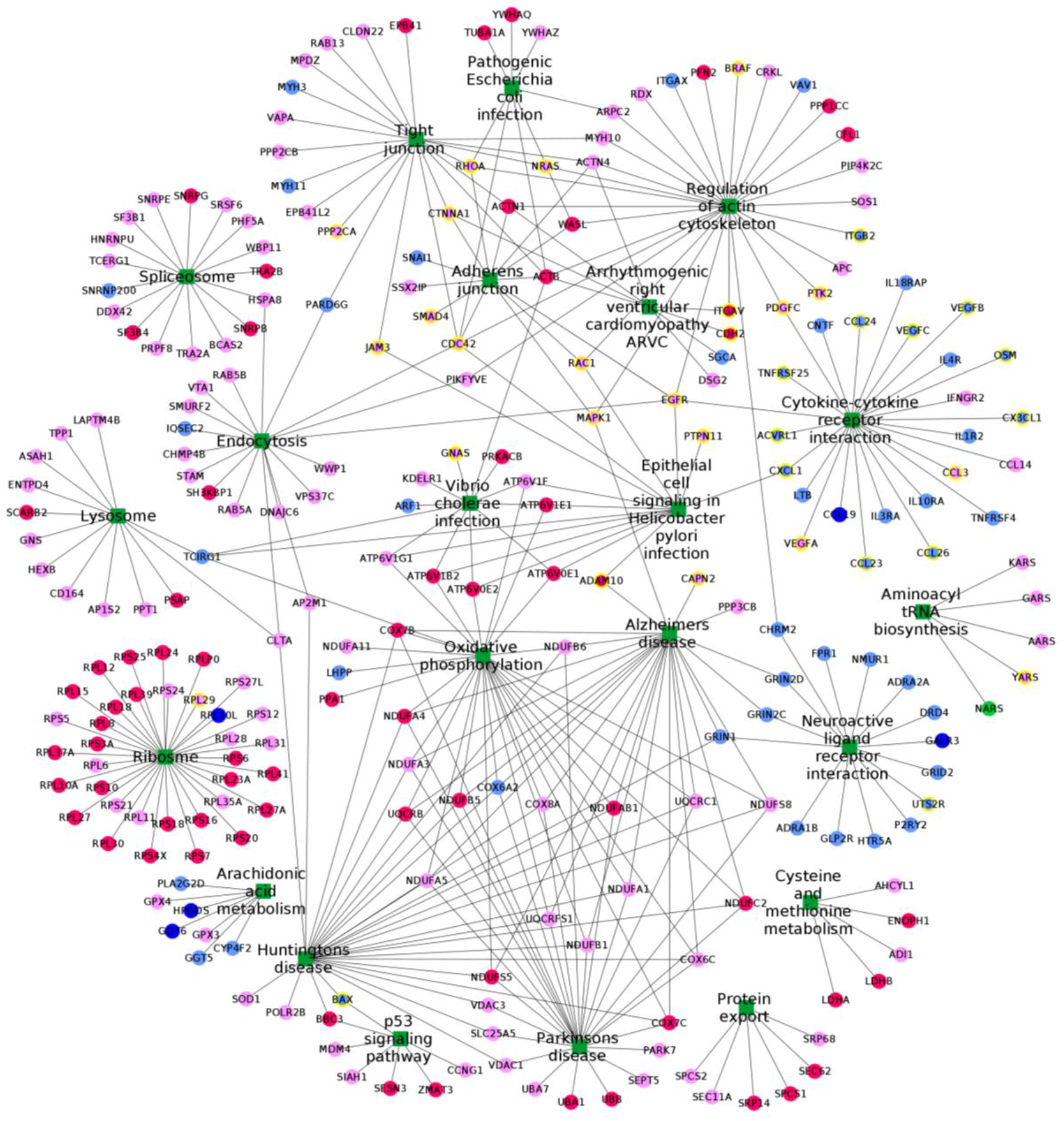
Dysregulated pathway interaction network for the deferentially expressed genes (DEGs). The green node represents the dysregulated pathways, pink represents DEGs with fold change between −3 and −2, red represents FC ≤ −3, pale blue represents FC between 2 and 3, dark blue represents FC ≥ 3. The round edged square shape of node represents pathways, circle represents DEGs, and yellow circle represents DEGs associated with angiogenesis GO.

The neuroactive ligand receptor interaction pathway was the highly dysregulated pathway with the normalized enrichment score (NES) of 3.15, and 15 DEGs were identified in this pathway. Of these, 11 DEGs were unique to this pathway and while, 4 DEGs GRIN1, CHRM2, GRIN2C and GRIN2D were connecting other pathways. Only UTS2R was found to be angiogenic-associated (Figure 2).

#### 3.2.2. Cytokine-cytokine receptor interaction pathway

Cytokine-cytokine receptor interaction pathway, next to the neuroactive ligand receptor interaction pathway, had enrichment score 2.55 (Table 1). We identified 25 DEGs (Figure 2 and S5) in this pathway. Among these, down-regulated VEGFA, CCL3, EGFR, PDGFC and up-regulated genes VEGFB, VEGFC, CX3CL1, ACVRL1, CCL23, CCL24, CCL26, OSM, and TNFRSF25 genes were found to be angiogenic-associated. Of interest, we identified that three of the angiogenic-associated genes PDGFC, CXCL1, and EGFR were also involved in the other pathways, as shown in Figures 2 and S5.

#### 3.2.3. Other pathways with DEGs of angiogenic association

In addition to highly enriched cytokine-cytokine receptor interaction and Neuroactive ligand receptor interaction pathways, we showed 13 other pathways with angiogenic-associated DEGs. Regulation of the actin cytoskeleton pathway has ten angiogenic DEGs; Adherens junction and Epithelial cell signaling in Helicobacter pylori infection have seven angiogenic DEGs; tight junction has six angiogenic DEGs. Three angiogenic DEGs were involved in Alzheimer’s disease and arrhythmogenic right ventricular cardiomyopathy ARVC pathways and 2 DEGs were in endocytosis and pathogenic Escherichia coli infection pathways. Five pathways had one DEG, namely, aminoacyl tRNA biosynthesis (YARS), Huntington’s disease (BAX), p53 signaling pathway (BAX), ribosome (RPL29) and Vibrio cholerae infection (GNAS).

The angiogenic-associated DEGs that were involved in two enriched pathways include ADAM10, BAX, CXCL1, ITGAV, JAM3, NRAS, and PDGFC. CTNNA1, MAPK1, and RAC1 were involved in three enriched pathways; RHOA in four pathways; EGFR and CDC42 were involved in more than five pathways. Despite the above, 92 angiogenic-associated DEGs were not involved in any of the enriched pathways. For instance, we identified the SEMA3B gene with high fold change of 5.154 and not involved in any of the enriched pathways.

#### 3.2.4. Angiogenic interactome of DEGs through protein-protein interaction (PPI)

In order to identify the interactome of angiogenic-associated DEGs, their protein-protein interaction (PPI) partners and pathways associated with them are connected as network and graphically illustrated in Figure 3. PPP2CA showed the highest connectivity, which is associated with the tight junction pathway, and it mostly connected to the ribosome pathway. Followed by PPP2CA, RHOA and EGFR showed more interacting partners and pathways.

**Figure 3.**
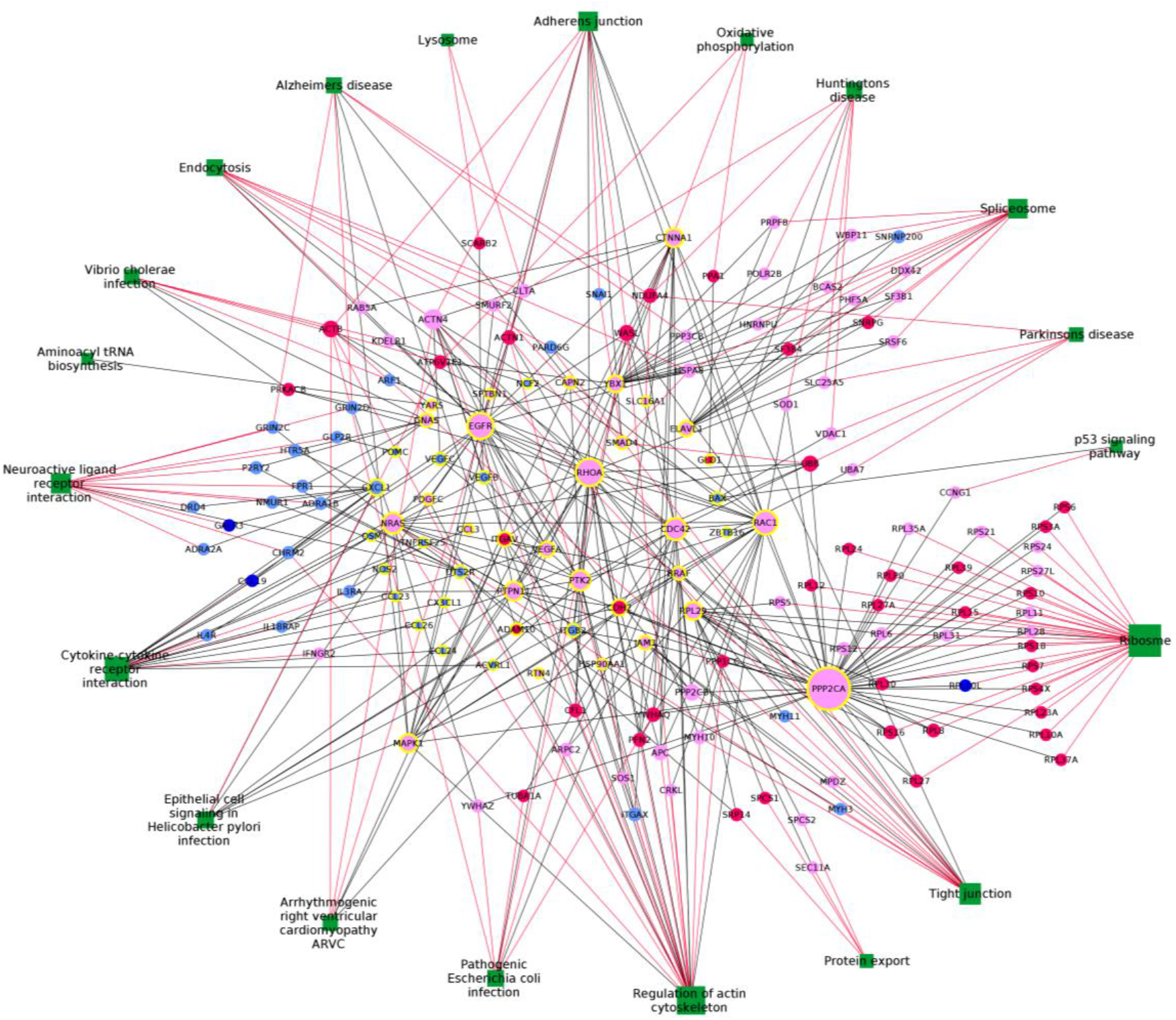
PPI network of core angiogenesis GO related differentially expressed genes and their PPI partners with the interaction of dysregulated pathways. The green node represents the dysregulated pathways, pink represents DEGs with fold change between −3 and −2, red represents FC ≤ −3, pale blue represents FC between 2 and 3, dark blue represents FC ≥ 3. The square shape of node represents pathways, circle represents DEGs, and yellow circle represents DEGs associated with angiogenesis GO. Edges red in colour represents the interaction between the pathway and the PPI partners of the DEGs which associated with angiogenesis. The size of the node corresponds to the degree i.e. number of edges linked to that node.

#### 3.2.5 Pathway specificity analysis

To examine the specificity of enriched pathways (Table 1), we performed a similar analysis using two sets of control samples, one from the retina and one from RPE/Choroid samples (Figure 1). Compare to enriched pathways obtained using all control samples; we identified that almost all the enriched pathways were enriched either using retina or RPE/Choroid or both except Cysteine and methionine metabolism and Vibrio cholerae infection. However, there are other pathways were enriched in the two control data sets that were not enriched in all control data as shown in Supplementary Table S1. Nevertheless, the enriched pathways, as shown in Table 1 is not affected by the two different retina and RPE/Choroid control data sets, not random, although they influence the pathway enrichment analysis.

#### 3.2.6 Disease-specific analysis

Furthermore, we performed disease-specific pathway enrichment analysis whether a single disease influences the angiogenic-associated pathways and genes. The heatmap of the dysregulated pathways with respect to disease specificity is illustrated in Figure 4. The highly enriched pathways Neuroactive ligand receptor interaction pathway and cytokine-cytokine receptor interaction pathway was not affected by the specific disease. The cytokine-cytokine receptor interaction pathway with highly angiogenic-associated DEGs was graphically illustrated as a heat map in Figure 5. Many angiogenic-associated genes (mentioned previously) showed similar patterns of fold change in all three retinal diseases. For example, VEGFA gene was downregulated in three all retinal diseases individually and all three together. We also performed the disease-specific comparative analysis using different control data set (Supplementary Table S2). When PDR samples compared with NPDR and DM samples, we observed the almost same pattern of dysregulated pathways. However, we did not observe similar pattern when we compared dry AMD with AMD samples.

**Figure 4.**
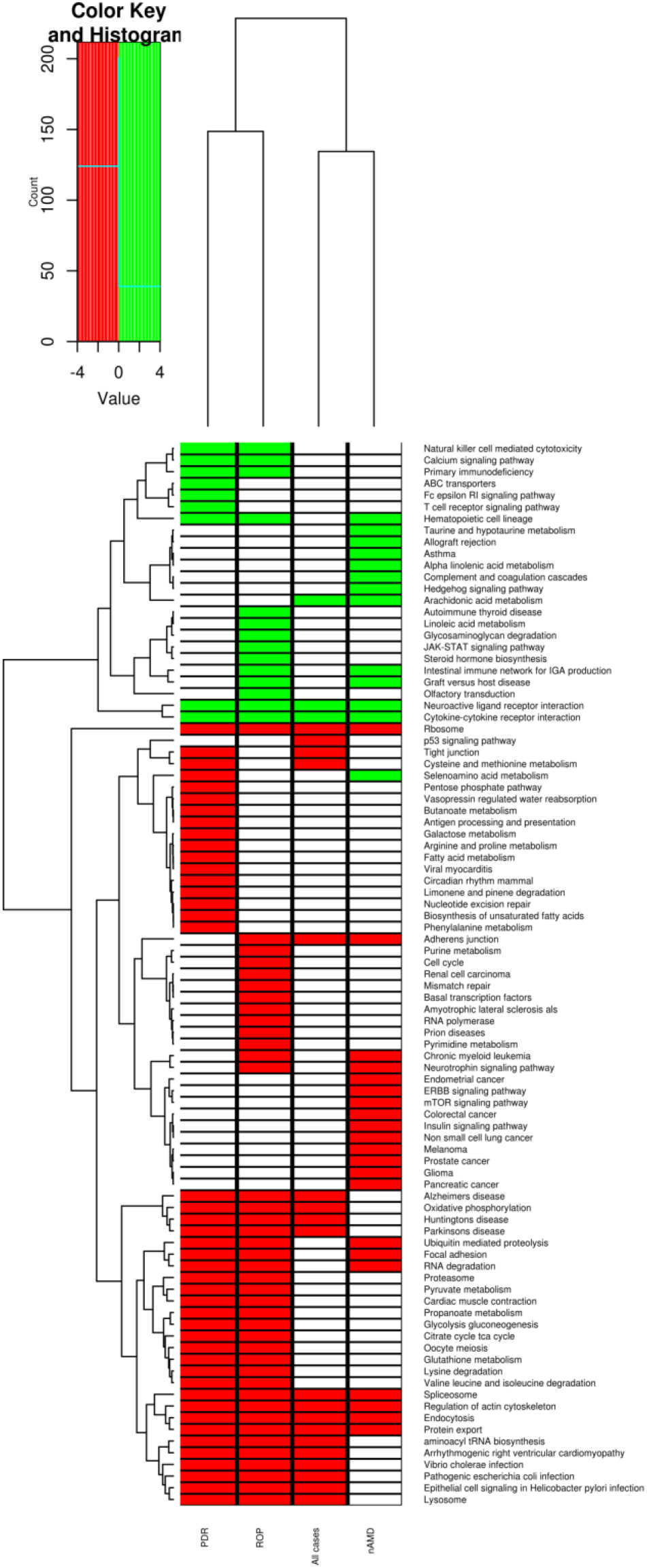
Clustering annotation with heat map of dysregulated pathways in retinal angiogenesis related diseases

**Figure 5.**
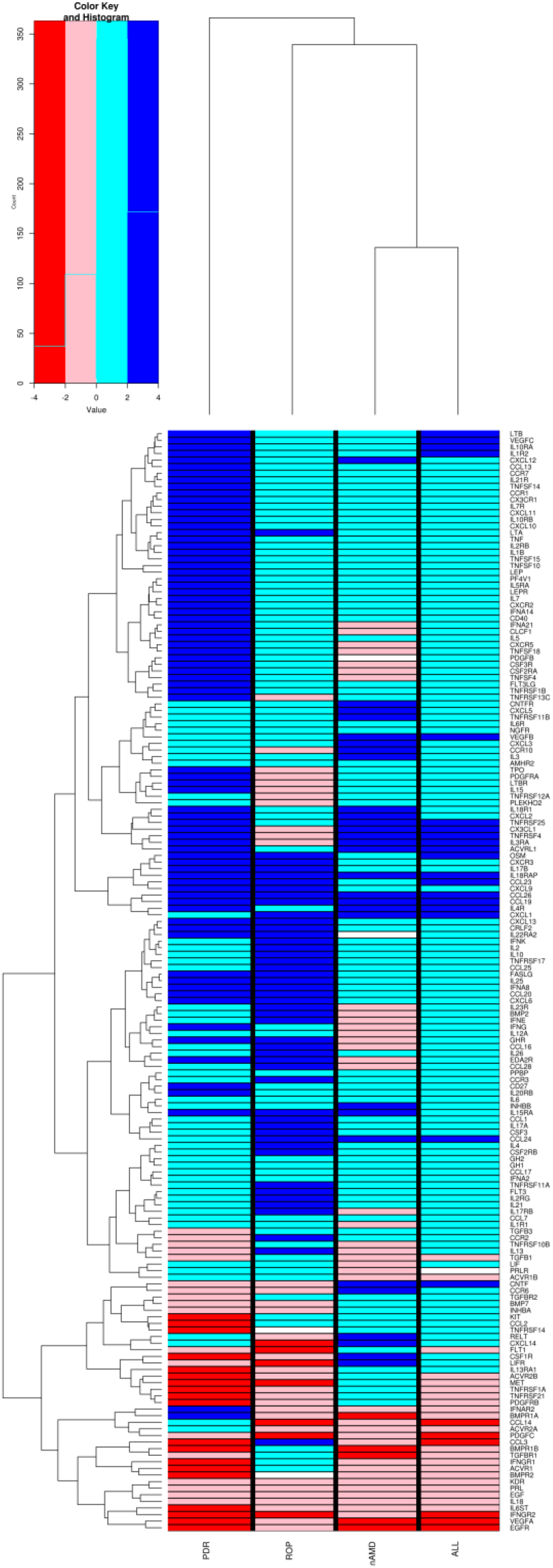
Heat map of DEGs in cytokine-cytokine receptor interaction pathway in retinal angiogenic related diseases

### 3.3. Pathway analysis using RNA-seq data

We further extended our analysis with the RNA-sequencing data set in order to show whether the pathway enrichment analysis is biased with microarray data. We compared nine PDR samples with 13 normal whole retina samples, and we identified 5816 upregulated and 2031 downregulated genes. The supplementary table 3 shows the dysregulated pathways with the NES score. The graft versus host disease pathway was enriched with the highest NES (4.15); however, it did not include the angiogenic DEGs. As expected, the Cytokine-cytokine receptor interaction pathway with NES of 3.35 is the most dysregulated in the retinal angiogenesis included with seventeen angiogenic-associated DEG. Among the DEGs, ACVRL1, CCL19, CCL23, CXCL1, IL10RA, IL18RAP, IL1R2, IL3RA, IL4R, OSM, TNFRSF25 and TNFRSF4 were observed with similar expression as that of the results from microarray data analysis. Notably, of the VEGF family, only VEGFC was observed with significant upregulation with 1.3 fold change.

## 4. Discussion

Neovascularization by dysregulation of angiogenic pathways is the major cause of vision loss in many retinal diseases. The purpose of this study is to identify dysregulated angiogenic pathways in addition to VEGF dependent angiogenic pathways and specific angiogenic factors involved in vision-threatening retinal diseases, include PDR, ROP and nAMD. We thought that the most feasible approach to this end would be to combine all the expression data from retinal diseases with angiogenesis, to identify common dysregulated angiogenic pathways. We showed that VEGF independent pathways are highly enriched by the differentially expressed genes (DEGs) of the retinal angiogenesis samples. The two highly enriched pathways are neuroactive ligand receptor interaction and cytokine-cytokine receptor interaction pathways (Figure 4). Among the several DEGs involved in neuroactive ligand receptor interaction pathway, though our angiome-network showed no angiogenic-associated genes, ADRA1B ^27^, CHRM2^28^, DRD4^29^, GALR3^30^, GLP2R^31^, GRIN2C^32^, GRIN2D^33^, HTR5A^34^, P2RY2^35^, UTS2R^36^ DEGs have been reported to be involved in angiogenesis. The neuroactive ligand receptor interaction pathway has been reported to be the most significant pathway for cancers and angiogenesis and is also closely connected with the poor prognosis of cancer^37–41^.

Based on the angiome-network with enriched DEGs, we report that the cytokine-cytokine receptor interaction pathway is the highly dysregulated angiogenic pathway enriched with fifteen angiogenic-associated DEGs (Supplementary Figure S5) followed by regulation of actin cytoskeleton, adherens junction and epithelial cell signaling in *Helicobacter pylori* infection. Jennifer et al. ^42^ experimentally identified novel VEGF-independent cytokines and their association with PDR, suggesting the association of cytokines in angiogenesis. In this study, we have shown that ten angiogenic-associated cytokines were significantly up-regulated in the dysregulated cytokine-cytokine receptor interaction pathway. Interestingly, VEGFA along with other angiogenic factors CCL3, EGFR and PDGFC are significantly down-regulated, suggesting that VEGF independent cytokines play a vital role in retinal angiogenesis. Apart from the ten up-regulated angiogenic-associated genes, with widespread literature survey, we account that CCL19^43^, IL18RAP^44^, IL1R2^45^, LTB^46^, IL10RA^47^, CNTF^48^ up-regulated genes may contribute to the dysregulation of angiogenesis. Therefore, we report that cytokine-cytokine receptor interaction pathway is the common dysregulated pathway in retinal angiogenesis as previously stated in the animal models of retinal angiogenesis^49^.

Of interest, to support that cytokine-cytokine receptor interaction pathway as common dysregulated angiogenic pathways in retinal diseases, we constructed a regulatory pathway (figure 6) with observed DEGs in this pathway and literature support. First, we exhibit that the VEGF pathway is downregulated by showing the downregulation of VEGFA gene expression (Figures 5 and 6) and upregulation of CCL19 with CCR7 in retinal angiogenesis. The CCL19 and CCR7 have been reported to regulate the VEGF pathway negatively^50–52^. Additionally, the DEGs CCL3^53^ and EGFR^54^ are reported to induce VEGFA expression, which is downregulated in this study. We observe the downregulation of other DEGs c-Met, RAS, ERK, Elk1 and STAT3 that could affect the VEGFA dependent angiogenesis^52^. The upregulated DEGs ITFRSF25, TNFRSF4, CCL23, CCL26, and CX3CL1 of cytokine-cytokine interaction pathway is reportedly activating PI3K/AKT pathway^55–59^, which in turn induces the angiogenesis through activating mTOR^54^ and NFKB pathway^60^. The upregulated DEGs IL18RAP and CXCL1has shown to modulates the NFKB and thus induces the angiogenesis. Furthermore, the upregulated DEG ACVRL1 is reported to promote endothelial cell proliferation by activating SMAD1^61^ and coexpression of IL12 with CCL19 reported to regulates the JAK/STAT signaling^62^, may help to mediate the angiogenesis indirectly^61–63^. We also observe that the other dysregulated pathways (Table 1) include aminoacyl tRNA biosynthesis, pathogenic Escherichia coli infection signaling, epithelial cell signaling in Helicobacter pylori infection and cysteine and methionine metabolism pathway, reported to trigger VEGF dependent angiogenic pathway^64–66^, are negatively enriched. We also support the fact that VEGF-independent angiogenic pathway in retinal angiogenesis using the PPI network. Most of the PPI partners of VEGFA namely EGF, FLT1, HIF1A, IGF1, KDR, PIK3CA, TGFB1, THBS1 are downregulated in this study. However, we have observed that the upregulation of VEGFB and VEGFC. However, comparing the expression VEGFB and VEGFC in each disease and results of RNA-seq analysis, we have identified that VEGFC expression is consistent, individually significant upregulation was observed in PDR (more than two FC).

**Figure 6.**
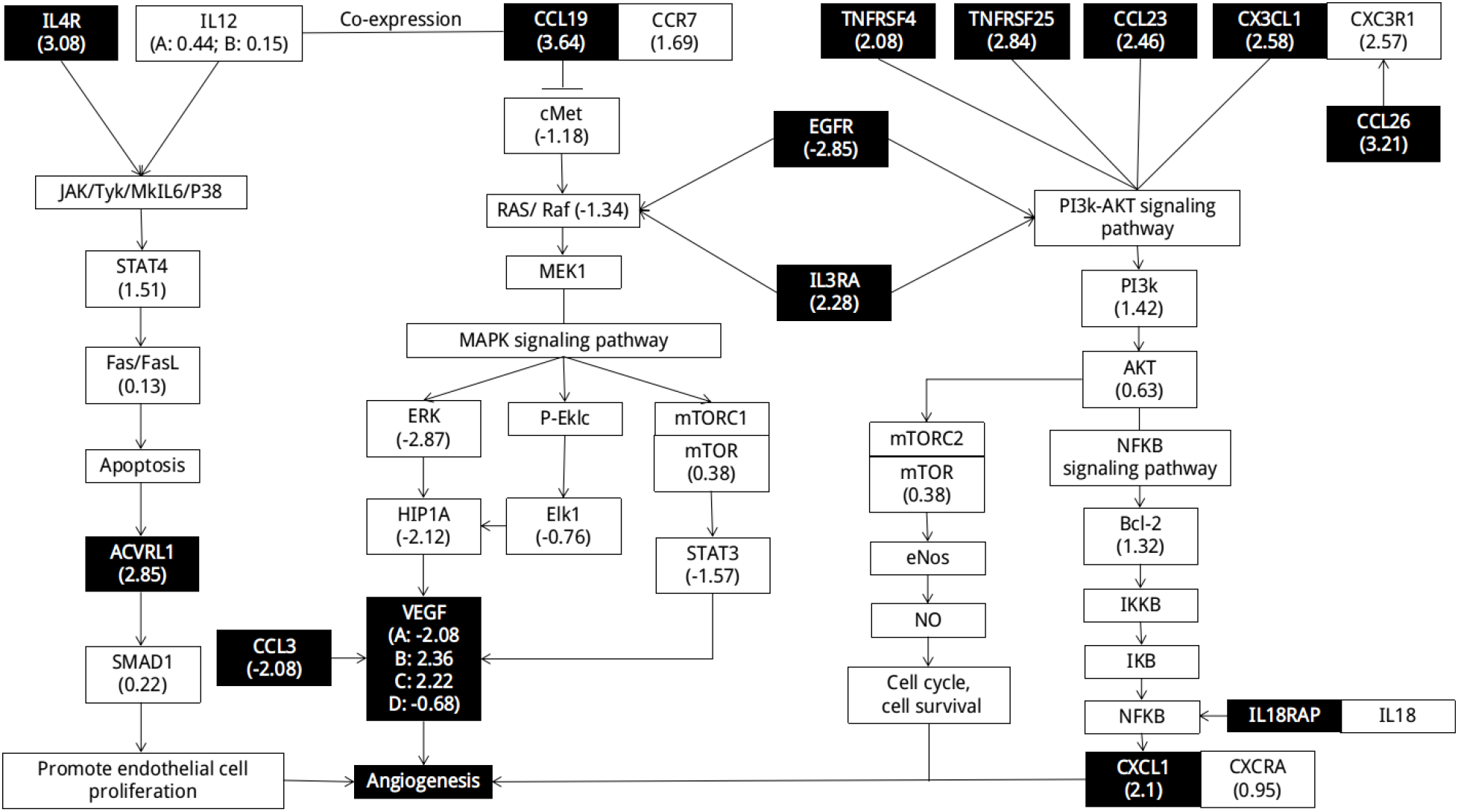
Regulatory role of DEGs in cytokine-cytokine receptor interaction pathway. Dark boxes indicate the DEGs identified in the retinal angiogenesis samples used in this study.

Further, to identify the specific angiogenic factors that are related to the cytokine-cytokine receptor interaction pathway, we have examined the DEGs in the disease-specific data. The fold change pattern of DEGs related to cytokine-cytokine receptor interaction pathway is following the same pattern. However, among DEGs, CCL19, CCL26, CCL24, CXCL1, ACVRL1, OSM, IL4R, CCL23and IL18RAP are significantly upregulated genes; while, VEGFA, EGFR, and IFNGR2 are downregulated genes in all three retinal diseases. The DEGs CCL14, CNTF, CX3CL1, IL10RA, IL15RA, IL1R2, IL3RA, LTB, PDGFC, TNFRSF25 and TNFRSF4 are upregulated genes in any of two retinal diseases. Further, when we compared the results with DEGs of cytokine-cytokine receptor interaction pathway enriched using RNA-Seq data, we have observed more than twelve DEGs that have similar expression viz ACVRL1, CCL19, CCL23, CXCL1, IL10RA, IL18RAP, IL1R2, IL3RA, IL4R, OSM, TNFRSF25 and TNFRSF4, suggesting their role in angiogenesis invariably. Furthermore, we have observed that CXCL1 is the major hub of upregulated DEGs based on the protein-protein interactions with the angiome-pathway network (figure 3), and thus it would be a potential target. In addition to CXCL1, based their angiogenic role (Figure 6), the PPI network (figure 3) and the RNA-seq data results, we highlight that VEGFC, ACVRL1, CCL23, CX3CL1 and TNFRSF25 are the notable targets.

In conclusion, the cytokine-cytokine receptor interaction pathway is highlighted as the common contributor for the retinal angiogenesis in PDR, ROP and nAMD through microarray metadata analysis. Further, several DEGs in the cytokine-cytokine receptor interaction pathway are identified as potential angiogenic factors that may cause the retinal angiogenesis. However, as only a few studies have been published related to the highlighted VEGF-independent retinal angiogenic pathways and genes, we hope that this work will lead future investigations into mechanisms of retinal angiogenesis, as well as identification of potential drug target.

## Acknowledgments

The authors are thankful to the Science and Engineering Research Board, Govt. of India (PDF/2016/001448), for financial support. The authors thank Prof. VR. Muthukkaruppan for the helpful discussion.

## Author Contributions

Conceptualization, U.S., methodology, U.S., writing—original draft preparation, U.S., Investigation, B.D., writing—review and editing, B.D., supervision, B.D.

