## Supplementary material for "Identification of Dysregulated Pathways and key genes in Human Retinal Angiogenesis using Microarray Metadata": Supplimentary Files

**Table S1. Dysregulated pathways of retinal angiogenesis samples compared to controls**

| S. No. | Pathway | Normalized enrichment score | | | | | |
|---|---|---|---|---|---|---|---|
| | | All cases Vs. All control | p-value | All cases Vs. control (Retina) | p-value | All cases Vs. Control (RPE/C) | p-value |
| 1 | Adherens junction | -2.04 | 0.00 | - | - | -1.95 | 0.00 |
| 2 | Alzheimers disease | -2.11 | 0.00 | -2.04 | 0.01 | -1.76 | 0.03 |
| 3 | aminoacyl tRNA biosynthesis | -1.57 | 0.04 | -1.62 | 0.03 | -1.67 | 0.02 |
| 4 | Arachidonic acid metabolism | 1.93 | 0.01 | 1.80 | 0.02 | 1.58 | 0.04 |
| 5 | Arrhythmogenic right ventricular cardiomyopathy | -1.89 | 0.01 | -1.67 | 0.04 | -2.35 | 0.00 |
| 6 | **Cysteine and methionine metabolism** | -1.67 | 0.03 | - | - | - | - |
| 7 | Cytokine-cytokine receptor interaction | 2.55 | 0.00 | 3.08 | 0.00 | - | - |
| 8 | Endocytosis | -1.77 | 0.01 | -1.70 | 0.02 | - | - |
| 9 | Epithelial cell signaling in Helicobacter pylori infection | -1.87 | 0.02 | - | - | -1.99 | 0.00 |
| 10 | Huntingtons disease | -2.49 | 0.00 | -1.81 | 0.01 | -1.94 | 0.01 |
| 11 | Lysosome | -1.93 | 0.00 | - | - | -2.21 | 0.00 |
| 12 | Neuroactive ligand receptor interaction | 3.10 | 0.00 | - | - | 2.81 | 0.00 |
| 13 | Oxidative phosphorylation | -2.58 | 0.00 | -2.31 | 0.00 | -2.36 | 0.00 |
| 14 | p53 signaling pathway | -1.89 | 0.01 | - | - | -1.84 | 0.02 |
| 15 | Parkinsons disease | -2.32 | 0.00 | -2.51 | 0.00 | -2.21 | 0.00 |
| 16 | Pathogenic escherichia coli infection | -1.65 | 0.03 | - | - | -1.78 | 0.01 |
| 17 | Protein export | -1.91 | 0.00 | -1.66 | 0.02 | -1.63 | 0.04 |
| 18 | Regulation of actin cytoskeleton | -1.87 | 0.01 | - | - | -2.33 | 0.00 |
| 19 | Rbosome | -4.05 | 0.00 | -2.61 | 0.00 | -3.83 | 0.00 |
| 20 | Spliceosome | -2.09 | 0.00 | -1.97 | 0.01 | -1.67 | 0.04 |
| 21 | Tight junction | -1.63 | 0.04 | - | - | -1.67 | 0.03 |
| 22 | **Vibrio cholerae infection** | -1.75 | 0.02 | - | - | - | - |
| 23 | Amyotrophic lateral sclerosis als | - | - | -1.71 | 0.03 | - | - |
| 24 | Antigen processing and presentation | - | - | -2.14 | 0.00 | -1.86 | 0.01 |
| 25 | Cytosolic dna sensing pathway | - | - | 1.56 | 0.05 | - | - |
| 26 | ERBB signaling pathway | - | - | -1.80 | 0.01 | -1.69 | 0.03 |
| 27 | Graft versus host disease | - | - | 1.68 | 0.02 | - | - |
| 28 | Hematopoietic cell lineage | - | - | 2.21 | 0.00 | - | - |
| 29 | Intestinal immune network for IGA production | - | - | 1.74 | 0.02 | - | - |
| 30 | JAK-STAT signaling pathway | - | - | 2.32 | 0.00 | - | - |

| S. No. | Pathway | Normalized enrichment score | | | | | |
|---|---|---|---|---|---|---|---|
| | | All cases Vs. All control | p-value | All cases Vs. control (Retina) | p-value | All cases Vs. Control (RPE/C) | p-value |
| 31 | Long term potentiation | - | - | -1.72 | 0.03 | - | - |
| 32 | Olfactory transduction | - | - | -1.82 | 0.01 | - | - |
| 33 | One carbon pool by folate | - | - | 1.57 | 0.04 | - | - |
| 34 | Primary immunodeficiency | - | - | 1.78 | 0.01 | - | - |
| 35 | Prion diseases | - | - | -1.66 | 0.02 | - | - |
| 36 | Progesterone mediated oocyte maturation | - | - | -1.69 | 0.04 | - | - |
| 37 | Pyruvate metabolism | - | - | -1.87 | 0.01 | - | - |
| 38 | RNA degradation | - | - | -1.81 | 0.02 | - | - |
| 39 | Small cell lung cancer | - | - | 1.87 | 0.00 | -1.99 | 0.00 |
| 40 | Ubiquitin mediated proteolysis | - | - | -1.90 | 0.00 | -1.59 | 0.04 |
| 41 | Biosynthesis of unsaturated fatty acids | - | - | - | - | -1.74 | 0.01 |
| 42 | Calcium signaling pathway | - | - | - | - | 2.10 | 0.00 |
| 43 | Chronic myeloid leukemia | - | - | - | - | -2.01 | 0.00 |
| 44 | Focal adhesion | - | - | - | - | -2.36 | 0.00 |
| 45 | Glioma | - | - | - | - | -1.69 | 0.04 |
| 46 | Melanoma | - | - | - | - | -2.04 | 0.00 |
| 47 | N glycan biosynthesis | - | - | - | - | -1.86 | 0.00 |
| 48 | Pancreatic cancer | - | - | - | - | -1.93 | 0.01 |
| 49 | Pathways in cancer | - | - | - | - | -1.73 | 0.02 |
| 50 | Peroxisome | - | - | - | - | -1.78 | 0.01 |
| 51 | Prostate cancer | - | - | - | - | -1.78 | 0.01 |
| 52 | VEGF signaling pathway | - | - | - | - | -1.57 | 0.05 |

### Table S2. Dysregulated pathways of PDR, ROP, nAMD compared to controls

| S. No. | Pathway | Normalized enrichment score | | | | | | | | | | | | | | | | | | | |
|---|---|---|---|---|---|---|---|---|---|---|---|---|---|---|---|---|---|---|---|---|---|
| | | PDR Vs. All control | p-value | PDR Vs. Control (Retina) | p-value | PDR Vs. Control (RPE/C) | p-value | ROP Vs. All control | p-value | nAMD Vs. All control | p-value | nMAD Vs. Control (Retina) | p-value | nAMD Vs. Control (RPE/C) | p-value | nAMD Vs. All dry AMD | p-value | nMAD Vs. Dry AMD (Retina) | p-value | nAMD Vs. Dry AMD (RPE/C) | p-value |
| 1 | Adherens junction | - | - | - | - | - | - | -1.84 | 0.01 | -1.64 | 0.03 | -2.38 | 0.00 | -2.14 | 0.00 | - | - | -1.78 | 0.02 | - | - |
| 2 | Alzheimers disease | -3.26 | 0.00 | -3.37 | 0.00 | -3.38 | 0.00 | -3.50 | 0.00 | - | - | - | - | - | - | - | - | - | - | - | - |
| 3 | Aminoacyl trna biosynthesis | -2.06 | 0.00 | - | - | - | - | -1.72 | 0.02 | - | - | - | - | - | - | - | - | - | - | - | - |
| 4 | Arachidonic acid metabolism | - | - | - | - | - | - | - | - | 1.90 | 0.01 | - | - | 2.13 | 0.00 | - | - | - | - | - | - |
| 5 | Arrhythmogenic right ventricular cardiomyopathy | -1.84 | 0.00 | - | - | - | - | -1.62 | 0.04 | - | - | - | - | -1.70 | 0.03 | - | - | - | - | - | - |
| 6 | Cysteine and methionine metabolism | -1.94 | 0.00 | -1.48 | 0.04 | - | - | - | - | - | - | - | - | - | - | - | - | - | - | - | - |

| S. No. | Pathway | Normalized enrichment score | | | | | | | | | | | | | | | | | | | |
|---|---|---|---|---|---|---|---|---|---|---|---|---|---|---|---|---|---|---|---|---|---|
| | | **PDR Vs. All control** | **p-value** | **PDR Vs. Control (Retina)** | **p-value** | **PDR Vs. Control (RPE/C)** | **p-value** | **ROP Vs. All control** | **p-value** | **nAMD Vs. All control** | **p-value** | **nMAD Vs. Control (Retina)** | **p-value** | **nAMD Vs. Control (RPE/C)** | **p-value** | **nAMD Vs. All dry AMD** | **p-value** | **nMAD Vs. Dry AMD (Retina)** | **p-value** | **nAMD Vs. Dry AMD (RPE/C)** | **p-value** |
| 7 | Cytokine cytokine receptor interaction | 2.98 | 0.00 | - | - | - | - | 3.80 | 0.00 | 2.14 | 0.00 | - | - | 2.13 | 0.00 | - | - | - | - | -1.38 | 0.00 |
| 8 | Endocytosis | -2.07 | 0.00 | - | - | - | - | -1.82 | 0.01 | -1.84 | 0.01 | - | - | -2.36 | 0.00 | - | - | - | - | - | - |
| 9 | Epithelial cell signaling in Helicobacter pylori infection | -2.24 | 0.00 | - | - | - | - | -1.78 | 0.02 | - | - | - | - | - | - | - | - | - | - | - | - |
| 10 | Huntingtons disease | -3.50 | 0.00 | -3.32 | 0.00 | -3.35 | 0.00 | -3.57 | 0.00 | - | - | - | - | - | - | - | - | - | - | - | - |
| 11 | Lysosome | -2.55 | 0.00 | - | - | - | - | -1.94 | 0.00 | - | - | - | - | -1.61 | 0.03 | - | - | - | - | - | - |
| 12 | Neuroactive | 3. | 0. | - | - | - | - | 4. | 0. | 1. | 0. | 1.6 | 0. | 2.0 | 0. | - | - | - | - | - | - |

| S. No. | Pathway | Normalized enrichment score | | | | | | | | | | | | | | | | | | | |
|---|---|---|---|---|---|---|---|---|---|---|---|---|---|---|---|---|---|---|---|---|---|
| | | PDR Vs. All control | p-value | PDR Vs. Control (Retina) | p-value | PDR Vs. Control (RPE/C) | p-value | ROP Vs. All control | p-value | nAMD Vs. All control | p-value | nMAD Vs. Control (Retina) | p-value | nAMD Vs. Control (RPE/C) | p-value | nAMD Vs. All dry AMD | p-value | nMAD Vs. Dry AMD (Retina) | p-value | nAMD Vs. Dry AMD (RPE/C) | p-value |
| | ligand receptor interaction | 36 | 00 | | | | | 39 | 00 | 84 | 02 | 0 | 04 | 2 | 00 | | | | | | |
| 13 | Oxidative phosphorylation | -4.03 | 0.00 | -3.50 | 0.00 | -3.53 | 0.00 | -3.64 | 0.00 | - | - | - | - | - | - | - | - | - | - | - | - |
| 14 | p53 signaling pathway | - | - | - | - | - | - | - | - | - | - | - | - | -2.19 | 0.00 | -2.43 | 0.00 | -3.05 | 0.00 | - | - |
| 15 | Parkinsons disease | -3.72 | 0.00 | -3.50 | 0.00 | -3.51 | 0.00 | -3.71 | 0.00 | - | - | - | - | -1.68 | 0.03 | - | - | - | - | - | - |
| 16 | Pathogenic escherichia coli infection | -2.26 | 0.00 | - | - | - | - | -2.43 | 0.00 | - | - | - | - | - | - | - | - | - | - | - | - |
| 17 | Protein export | - | 0. | - | - | - | - | - | 0. | - | 0. | - | 0. | - | 0. | - | - | - | - | - | - |

| S. No. | Pathway | Normalized enrichment score | | | | | | | | | | | | | | | | | | | |
|---|---|---|---|---|---|---|---|---|---|---|---|---|---|---|---|---|---|---|---|---|---|
| | | PDR Vs. All control | p-value | PDR Vs. Control (Retina) | p-value | PDR Vs. Control (RPE/C) | p-value | ROP Vs. All control | p-value | nAMD Vs. All control | p-value | nMAD Vs. Control (Retina) | p-value | nAMD Vs. Control (RPE/C) | p-value | nAMD Vs. All dry AMD | p-value | nMADD Vs. Dry AMD (Retina) | p-value | nAMD Vs. Dry AMD (RPE/C) | p-value |
| | | 2.01 | 00 | | | | | 2.05 | 00 | 2.03 | 00 | 1.86 | 00 | 2.14 | 00 | | | | | | |
| 18 | Regulation of actin cytoskeleton | -1.62 | 0.03 | - | - | - | - | -1.71 | 0.02 | -1.62 | 0.04 | -1.72 | 0.02 | - | - | -1.77 | 0.01 | -2.81 | 0.00 | - | - |
| 19 | Ribosome | -2.83 | 0.00 | - | - | - | - | -4.25 | 0.00 | -4.56 | 0.00 | -3.28 | 0.00 | -4.46 | 0.00 | - | - | - | - | - | - |
| 20 | Spliceosome | -2.79 | 0.00 | - | - | - | - | -2.78 | 0.00 | -1.67 | 0.03 | -1.65 | 0.05 | -1.83 | 0.02 | - | - | - | - | - | - |
| 21 | Tight junction | -1.61 | 0.02 | - | - | - | - | - | - | - | - | - | - | - | - | - | - | - | - | - | - |
| 22 | Vibrio cholerae infection | -3.01 | 0.00 | - | - | - | - | -1.95 | 0.01 | - | - | 1.65 | 0.02 | - | - | - | - | - | - | - | - |
| 23 | Amyotrophic | - | - | - | - | - | - | - | 0. | - | - | 2.0 | 0. | - | - | 1. | 0. | 1.8 | 0. | - | - |

| S. No. | Pathway | Normalized enrichment score | | | | | | | | | | | | | | | | | | | |
|---|---|---|---|---|---|---|---|---|---|---|---|---|---|---|---|---|---|---|---|---|---|
| | | PDR Vs. All control | p-value | PDR Vs. Control (Retina) | p-value | PDR Vs. Control (RPE/C) | p-value | ROP Vs. All control | p-value | nAMD Vs. All control | p-value | nMAD Vs. Control (Retina) | p-value | nAMD Vs. Control (RPE/C) | p-value | nAMD Vs. All dry AMD | p-value | nMAD Vs. Dry AMD (Retina) | p-value | nAMD Vs. Dry AMD (RPE/C) | p-value |
| | lateral sclerosis als | | | | | | | 1. 81 | 01 | | | 5 | 00 | | | 82 | 02 | 8 | 01 | | |
| 24 | Antigen processing and presentation | -2.12 | 0.00 | - | - | - | - | - | - | - | - | - | - | - | - | - | - | - | - | - | - |
| 25 | Cytosolic dna sensing pathway | - | - | - | - | - | - | - | - | - | - | - | - | - | - | - | - | - | - | - | - |
| 26 | ERBB signaling pathway | - | - | - | - | - | - | - | - | -2.04 | 0.00 | -2.01 | 0.00 | -1.65 | 0.04 | - | - | - | - | - | - |
| 27 | Graft versus host disease | - | - | - | - | - | - | 1.66 | 0.03 | 1.61 | 0.04 | - | - | 2.19 | 0.00 | - | - | - | - | - | - |
| 28 | Hematopoietic cell lineage | 3.15 | 0.00 | - | - | - | - | 1.63 | 0.05 | 1.95 | 0.01 | - | - | 2.29 | 0.00 | - | - | - | - | - | - |
| 29 | Intestinal immune network for iga | - | - | - | - | - | - | 2.02 | 0.00 | 1.89 | 0.01 | - | - | 2.28 | 0.00 | - | - | - | - | - | - |

| S. No. | Pathway | Normalized enrichment score | | | | | | | | | | | | | | | | | | | |
|---|---|---|---|---|---|---|---|---|---|---|---|---|---|---|---|---|---|---|---|---|---|
| | | PDR Vs. All control | p-value | PDR Vs. Control (Retina) | p-value | PDR Vs. Control (RPE/C) | p-value | ROP Vs. All control | p-value | nAMD Vs. All control | p-value | nMAD Vs. Control (Retina) | p-value | nAMD Vs. Control (RPE/C) | p-value | nAMD Vs. All dry AMD | p-value | nMAD Vs. Dry AMD (Retina) | p-value | nAMD Vs. Dry AMD (RPE/C) | p-value |
| | production | | | | | | | | | | | | | | | | | | | | |
| 30 | JAK stat signaling pathway | - | - | - | - | - | - | 2.07 | 0.01 | - | - | - | - | - | - | - | - | -1.63 | 0.04 | - | - |
| 31 | Long term potentiation | - | - | - | - | - | - | - | - | - | - | - | - | - | - | - | - | 1.63 | 0.02 | - | - |
| 32 | Olfactory transduction | - | - | - | - | - | - | 4.54 | 0.00 | - | - | - | - | - | - | - | - | 1.82 | 0.00 | - | - |
| 33 | One carbon pool by folate | - | - | - | - | - | - | - | - | - | - | - | - | - | - | - | - | - | - | - | - |
| 34 | Primary immunodeficiency | 2.25 | 0.00 | - | - | - | - | 2.20 | 0.01 | - | - | - | - | - | - | - | - | - | - | - | - |
| 35 | Prion diseases | - | - | - | - | - | - | -1.88 | 0.01 | - | - | - | - | - | - | - | - | - | - | - | - |
| 36 | Progesterone mediated oocyte | - | - | - | - | - | - | - | - | - | - | - | - | - | - | - | - | - | - | - | - |

| S. No. | Pathway | Normalized enrichment score | | | | | | | | | | | | | | | | | |
|---|---|---|---|---|---|---|---|---|---|---|---|---|---|---|---|---|---|---|---|
| | | PDR Vs. All control | p-value | PDR Vs. Control (Retina) | p-value | PDR Vs. Control (RPE/C) | p-value | ROP Vs. All control | p-value | nAMD Vs. All control | p-value | nMAD Vs. Control (Retina) | p-value | nAMD Vs. Control (RPE/C) | p-value | nAMD Vs. All dry AMD | p-value | nMAD Vs. Dry AMD (Retina) | p-value | nAMD Vs. Dry AMD (RPE/C) | p-value |
| | maturation | | | | | | | | | | | | | | | | | | | | |
| 37 | Pyruvate metabolism | -2.65 | 0.00 | - | - | - | - | -2.12 | 0.00 | - | - | - | - | - | - | - | - | - | - | - | - |
| 38 | RNA degradation | -1.94 | 0.01 | - | - | - | - | -1.96 | 0.01 | -1.73 | 0.02 | - | - | -2.16 | 0.00 | - | - | - | - | - | - |
| 39 | Small cell lung cancer | - | - | - | - | - | - | - | - | - | - | - | - | -2.35 | 0.00 | - | - | -2.74 | 0.00 | - | - |
| 40 | Ubiquitin mediated proteolysis | -1.92 | 0.00 | - | - | - | - | -2.55 | 0.00 | -1.67 | 0.03 | -1.80 | 0.02 | -1.94 | 0.00 | - | - | - | - | - | - |
| 41 | Biosynthesis of | - | 0. | - | - | - | - | - | - | - | - | - | - | - | - | - | - | - | - | - | - |

| S. No. | Pathway | Normalized enrichment score | | | | | | | | | | | | | | | | | | | |
|---|---|---|---|---|---|---|---|---|---|---|---|---|---|---|---|---|---|---|---|---|---|
| | | PDR Vs. All control | p-value | PDR Vs. Control (Retina) | p-value | PDR Vs. Control (RPE/C) | p-value | ROP Vs. All control | p-value | nAMD Vs. All control | p-value | nMAD Vs. Control (Retina) | p-value | nAMD Vs. Control (RPE/C) | p-value | nAMD Vs. All dry AMD | p-value | nMAD Vs. Dry AMD (Retina) | p-value | nAMD Vs. Dry AMD (RPE/C) | p-value |
| | unsaturated fatty acids | 1.77 | 01 | | | | | | | | | | | | | | | | | | |
| 42 | Calcium signaling pathway | 1.70 | 0.01 | - | - | - | - | 2.04 | 0.00 | - | - | - | - | - | - | - | - | - | - | - | - |
| 43 | Chronic myeloid leukemia | - | - | - | - | - | - | -1.61 | 0.05 | -2.08 | 0.00 | -2.28 | 0.00 | -2.30 | 0.00 | - | - | -1.90 | 0.01 | - | - |
| 44 | Focal adhesion | -1.56 | 0.03 | - | - | - | - | -1.89 | 0.00 | -1.80 | 0.01 | - | - | -1.67 | 0.02 | - | - | -3.15 | 0.00 | - | - |
| 45 | Glioma | - | - | - | - | - | - | - | - | -1.92 | 0.01 | - | - | -1.55 | 0.05 | - | - | - | - | - | - |
| 46 | Melanoma | - | - | - | - | - | - | - | - | -1.83 | 0.01 | - | - | -1.87 | 0.02 | - | - | - | - | - | - |
| 47 | N glycan | - | - | - | - | - | - | - | - | - | - | - | - | - | - | - | - | - | - | - | - |

| S. No. | Pathway | Normalized enrichment score | | | | | | | | | | | | | | | | | | | |
|---|---|---|---|---|---|---|---|---|---|---|---|---|---|---|---|---|---|---|---|---|---|
| | | PDR Vs. All control | p-value | PDR Vs. Control (Retina) | p-value | PDR Vs. Control (RPE/C) | p-value | ROP Vs. All control | p-value | nAMD Vs. All control | p-value | nMAD Vs. Control (Retina) | p-value | nAMD Vs. Control (RPE/C) | p-value | nAMD Vs. All dry AMD | p-value | nMAD Vs. Dry AMD (Retina) | p-value | nAMD Vs. Dry AMD (RPE/C) | p-value |
| | biosynthesis | | | | | | | | | | | | | | | | | | | | |
| 48 | Pancreatic cancer | - | - | - | - | - | - | - | - | -1.94 | 0.01 | -1.91 | 0.01 | -2.30 | 0.00 | - | - | -1.89 | 0.00 | - | - |
| 49 | Pathways in cancer | - | - | - | - | - | - | - | - | - | - | - | - | -1.62 | 0.04 | - | - | -2.27 | 0.00 | - | - |
| 50 | Peroxisome | - | - | - | - | - | - | - | - | - | - | - | - | -1.95 | 0.00 | - | - | - | - | - | - |
| 51 | Prostate cancer | - | - | - | - | - | - | - | - | -1.87 | 0.01 | - | - | -1.75 | 0.02 | - | - | - | - | - | - |
| 52 | VEGF signaling pathway | - | - | - | - | - | - | - | - | - | - | - | - | - | - | - | - | - | - | - | - |
| 53 | ABC transporters | 1.92 | 0.01 | - | - | - | - | - | - | - | - | - | - | - | - | - | - | - | - | - | - |
| 54 | Arginine and | - | 0. | - | - | - | - | - | - | - | - | - | - | - | - | - | - | - | - | - | - |

| S. No. | Pathway | Normalized enrichment score | | | | | | | | | | | | | | | | | | | |
|---|---|---|---|---|---|---|---|---|---|---|---|---|---|---|---|---|---|---|---|---|---|
| | | PDR Vs. All control | p-value | PDR Vs. Control (Retina) | p-value | PDR Vs. Control (RPE/C) | p-value | ROP Vs. All control | p-value | nAMD Vs. All control | p-value | nMAD Vs. Control (Retina) | p-value | nAMD Vs. Control (RPE/C) | p-value | nAMD Vs. All dry AMD | p-value | nMAD Vs. Dry AMD (Retina) | p-value | nAMD Vs. Dry AMD (RPE/C) | p-value |
| | proline metabolism | 1.65 | 03 | | | | | | | | | | | | | | | | | | |
| 55 | Butanoate metabolism | -2.11 | 0.00 | - | - | - | - | - | - | - | - | - | - | - | - | - | - | - | - | - | - |
| 56 | Cardiac muscle contraction | -2.42 | 0.00 | -2.74 | 0.00 | -2.71 | 0.00 | -2.60 | 0.00 | - | - | - | - | - | - | 1.66 | 0.01 | - | - | - | - |
| 57 | Circadian rhythm mammal | -1.86 | 0.00 | - | - | - | - | - | - | - | - | - | - | - | - | - | - | - | - | - | - |
| 58 | Citrate cycle tca cycle | -1.97 | 0.00 | - | - | - | - | -2.26 | 0.00 | - | - | - | - | - | - | - | - | - | - | - | - |
| 59 | Fatty acid metabolism | -1.61 | 0.05 | - | - | - | - | - | - | - | - | - | - | - | - | - | - | - | - | - | - |
| 60 | FC epsilon RI | 1. | 0. | - | - | - | - | - | - | - | - | - | - | - | - | - | - | - | - | - | - |

| S. No. | Pathway | Normalized enrichment score | | | | | | | | | | | | | | | | | | | |
|---|---|---|---|---|---|---|---|---|---|---|---|---|---|---|---|---|---|---|---|---|---|
| | | PDR Vs. All control | p-value | PDR Vs. Control (Retina) | p-value | PDR Vs. Control (RPE/C) | p-value | ROP Vs. All control | p-value | nAMD Vs. All control | p-value | nMAD Vs. Control (Retina) | p-value | nAMD Vs. Control (RPE/C) | p-value | nAMD Vs. All dry AMD | p-value | nMAD Vs. Dry AMD (Retina) | p-value | nAMD Vs. Dry AMD (RPE/C) | p-value |
| | signaling pathway | 84 | 01 | | | | | | | | | | | | | | | | | | |
| 61 | Galactose metabolism | -1.59 | 0.04 | - | - | - | - | - | - | - | - | - | - | - | - | - | - | - | - | - | - |
| 62 | Glutathione metabolism | -1.83 | 0.01 | - | - | - | - | -1.58 | 0.04 | - | - | - | - | - | - | - | - | - | - | - | - |
| 63 | Glycolysis gluconeogenesis | -2.44 | 0.00 | - | - | - | - | -1.75 | 0.02 | - | - | - | - | -2.55 | 0.00 | - | - | - | - | - | - |
| 64 | Limonene and pinene degradation | -1.69 | 0.01 | - | - | - | - | - | - | - | - | - | - | - | - | - | - | - | - | - | - |
| 65 | Lysine degradation | -1.95 | 0.00 | - | - | - | - | -1.73 | 0.02 | - | - | - | - | - | - | - | - | - | - | - | - |
| 66 | Natural killer | 1. | 0. | - | - | - | - | 2. | 0. | - | - | - | - | - | - | - | - | - | - | - | - |

| S. No. | Pathway | Normalized enrichment score | | | | | | | | | | | | | | | | | | | |
|---|---|---|---|---|---|---|---|---|---|---|---|---|---|---|---|---|---|---|---|---|---|
| | | PDR Vs. All control | p-value | PDR Vs. Control (Retina) | p-value | PDR Vs. Control (RPE/C) | p-value | ROP Vs. All control | p-value | nAMD Vs. All control | p-value | nMAD Vs. Control (Retina) | p-value | nAMD Vs. Control (RPE/C) | p-value | nAMD Vs. All dry AMD | p-value | nMAD Vs. Dry AMD (Retina) | p-value | nAMD Vs. Dry AMD (RPE/C) | p-value |
| | cell mediated cytotoxicity | 76 | 01 | | | | | 06 | 00 | | | | | | | | | | | | |
| 67 | Nucleotide excision repair | -1.76 | 0.02 | - | - | - | - | - | - | - | - | - | - | - | - | - | - | - | - | - | - |
| 68 | Oocyte meiosis | -1.62 | 0.03 | - | - | - | - | -1.88 | 0.01 | - | - | - | - | - | - | - | - | - | - | - | - |
| 69 | Pentose phosphate pathway | -2.16 | 0.00 | - | - | - | - | - | - | - | - | - | - | - | - | - | - | - | - | - | - |
| 70 | Phenylalanine metabolism | -1.77 | 0.00 | - | - | - | - | - | - | - | - | - | - | - | - | - | - | - | - | - | - |
| 71 | Propanoate metabolism | -2.22 | 0.00 | - | - | - | - | -1.61 | 0.05 | - | - | - | - | - | - | - | - | - | - | - | - |
| 72 | Proteasome | - | 0. | - | - | - | - | - | 0. | - | - | - | - | - | - | - | - | - | - | - | - |

| S. No. | Pathway | Normalized enrichment score | | | | | | | | | | | | | | | | | | | |
|---|---|---|---|---|---|---|---|---|---|---|---|---|---|---|---|---|---|---|---|---|---|
| | | PDR Vs. All control | p-value | PDR Vs. Control (Retina) | p-value | PDR Vs. Control (RPE/C) | p-value | ROP Vs. All control | p-value | nAMD Vs. All control | p-value | nMAD Vs. Control (Retina) | p-value | nAMD Vs. Control (RPE/C) | p-value | nAMD Vs. All dry AMD | p-value | nMAD Vs. Dry AMD (Retina) | p-value | nAMD Vs. Dry AMD (RPE/C) | p-value |
| | | 2.94 | 00 | | | | | 2.90 | 00 | | | | | | | | | | | | |
| 73 | Selenoamino acid metabolism | -1.67 | 0.02 | - | - | - | - | - | - | 1.61 | 0.03 | - | - | 1.71 | 0.01 | - | - | - | - | - | - |
| 74 | T cell receptor signaling pathway | 1.51 | 0.04 | - | - | - | - | - | - | - | - | - | - | - | - | - | - | - | - | - | - |
| 75 | Valine leucine and isoleucine degradation | -2.02 | 0.00 | - | - | - | - | -1.84 | 0.01 | - | - | - | - | - | - | - | - | - | - | - | - |
| 76 | Vasopressin regulated water reabsorption | -2.07 | 0.00 | - | - | - | - | - | - | - | - | - | - | - | - | - | - | - | - | - | - |
| 77 | Viral myocarditis | -1.63 | 0.03 | - | - | - | - | - | - | - | - | - | - | - | - | - | - | - | - | - | - |
| 78 | Adipocytokine | - | - | 1.6 | 0. | 1.6 | 0. | - | - | - | - | - | - | - | - | - | - | - | - | - | - |

| S. No. | Pathway | Normalized enrichment score | | | | | | | | | | | | | | | | | | | |
|---|---|---|---|---|---|---|---|---|---|---|---|---|---|---|---|---|---|---|---|---|---|
| | | PDR Vs. All control | p-value | PDR Vs. Control (Retina) | p-value | PDR Vs. Control (RPE/C) | p-value | ROP Vs. All control | p-value | nAMD Vs. All control | p-value | nMAD Vs. Control (Retina) | p-value | nAMD Vs. Control (RPE/C) | p-value | nAMD Vs. All dry AMD | p-value | nMAD Vs. Dry AMD (Retina) | p-value | nAMD Vs. Dry AMD (RPE/C) | p-value |
| | signaling pathway | | | 3 | 02 | 6 | 02 | | | | | | | | | | | | | | |
| 79 | PPARr signaling pathway | - | - | 2.60 | 0.00 | 2.37 | 0.00 | - | - | - | - | - | - | - | - | - | - | - | - | - | - |
| 80 | Apoptosis | - | - | - | - | 1.78 | 0.01 | - | - | - | - | - | - | - | - | - | - | - | - | - | - |
| 81 | Autoimmune thyroid disease | - | - | - | - | - | - | 1.65 | 0.03 | - | - | - | - | 2.15 | 0.00 | - | - | - | - | - | - |
| 82 | Basal transcription factors | - | - | - | - | - | - | -1.70 | 0.03 | - | - | - | - | - | - | - | - | - | - | - | - |
| 83 | Cell cycle | - | - | - | - | - | - | -2.23 | 0.00 | - | - | - | - | -2.34 | 0.00 | - | - | -2.13 | 0.00 | - | - |
| 84 | Glycosaminoglycan degradation | - | - | - | - | - | - | 1.51 | 0.04 | - | - | - | - | - | - | - | - | - | - | - | - |
| 85 | Linoleic acid metabolism | - | - | - | - | - | - | 1.62 | 0.03 | - | - | - | - | 1.66 | 0.02 | - | - | - | - | - | - |

| S. No. | Pathway | Normalized enrichment score | | | | | | | | | | | | | | | | | | | |
|---|---|---|---|---|---|---|---|---|---|---|---|---|---|---|---|---|---|---|---|---|---|
| | | PDR Vs. All control | p-value | PDR Vs. Control (Retina) | p-value | PDR Vs. Control (RPE/C) | p-value | ROP Vs. All control | p-value | nAMD Vs. All control | p-value | nMAD Vs. Control (Retina) | p-value | nAMD Vs. Control (RPE/C) | p-value | nAMD Vs. All dry AMD | p-value | nMAD Vs. Dry AMD (Retina) | p-value | nAMD Vs. Dry AMD (RPE/C) | p-value |
| 86 | Mismatch repair | - | - | - | - | - | - | -1.67 | 0.02 | - | - | - | - | - | - | - | - | - | - | - | - |
| 87 | Neurotrophin signaling pathway | - | - | - | - | - | - | -1.81 | 0.02 | -1.87 | 0.01 | - | - | - | - | - | - | - | - | - | - |
| 88 | Purine metabolism | - | - | - | - | - | - | -2.07 | 0.00 | - | - | - | - | - | - | - | - | - | - | - | - |
| 89 | Pyrimidine metabolism | - | - | - | - | - | - | -1.92 | 0.01 | - | - | - | - | - | - | - | - | - | - | - | - |
| 90 | Renal cell carcinoma | - | - | - | - | - | - | -1.65 | 0.04 | - | - | - | - | -1.70 | 0.03 | - | - | - | - | - | - |
| 91 | RNA polymerase | - | - | - | - | - | - | -1.96 | 0.00 | - | - | - | - | - | - | - | - | - | - | - | - |

| S. No. | Pathway | Normalized enrichment score | | | | | | | | | | | | | | | | | | | |
|---|---|---|---|---|---|---|---|---|---|---|---|---|---|---|---|---|---|---|---|---|---|
| | | PDR Vs. All control | p-value | PDR Vs. Control (Retina) | p-value | PDR Vs. Control (RPE/C) | p-value | ROP Vs. All control | p-value | nAMD Vs. All control | p-value | nMAD Vs. Control (Retina) | p-value | nAMD Vs. Control (RPE/C) | p-value | nAMD Vs. All dry AMD | p-value | nMAD Vs. Dry AMD (Retina) | p-value | nAMD Vs. Dry AMD (RPE/C) | p-value |
| 92 | Steroid hormone biosynthesis | - | - | - | - | - | - | 1.94 | 0.01 | - | - | - | - | - | - | - | - | - | - | - | - |
| 93 | Allograft rejection | - | - | - | - | - | - | - | - | 1.60 | 0.04 | - | - | 2.17 | 0.00 | - | - | - | - | - | - |
| 94 | Alpha linolenic acid metabolism | - | - | - | - | - | - | - | - | 1.53 | 0.05 | - | - | 1.88 | 0.01 | - | - | - | - | - | - |
| 95 | Asthma | - | - | - | - | - | - | - | - | 1.55 | 0.05 | - | - | 2.01 | 0.00 | - | - | - | - | - | - |
| 96 | Colorectal cancer | - | - | - | - | - | - | - | - | -1.74 | 0.01 | - | - | -1.87 | 0.00 | - | - | - | - | - | - |
| 97 | Complement and coagulation cascades | - | - | - | - | - | - | - | - | 1.82 | 0.03 | - | - | 2.28 | 0.00 | - | - | - | - | - | - |
| 98 | Endometrial cancer | - | - | - | - | - | - | - | - | -2.40 | 0.00 | -1.79 | 0.01 | -2.19 | 0.00 | - | - | - | - | - | - |
| 99 | Hedgehog | - | - | - | - | - | - | - | - | 1. | 0. | - | - | - | - | - | - | - | - | - | - |

| S. No. | Pathway | Normalized enrichment score | | | | | | | | | | | | | | | | | | | |
|---|---|---|---|---|---|---|---|---|---|---|---|---|---|---|---|---|---|---|---|---|---|
| | | PDR Vs. All control | p-value | PDR Vs. Control (Retina) | p-value | PDR Vs. Control (RPE/C) | p-value | ROP Vs. All control | p-value | nAMD Vs. All control | p-value | nMAD Vs. Control (Retina) | p-value | nAMD Vs. Control (RPE/C) | p-value | nAMD Vs. All dry AMD | p-value | nMAD Vs. Dry AMD (Retina) | p-value | nAMD Vs. Dry AMD (RPE/C) | p-value |
| | signaling pathway | | | | | | | | | 79 | 01 | | | | | | | | | | |
| 100 | Insulin signaling pathway | - | - | - | - | - | - | - | - | -1.67 | 0.03 | - | - | - | - | - | - | - | - | - | - |
| 101 | mTOR signaling pathway | - | - | - | - | - | - | - | - | -2.15 | 0.00 | - | - | -1.79 | 0.01 | - | - | - | - | - | - |
| 102 | Non small cell lung cancer | - | - | - | - | - | - | - | - | -1.68 | 0.04 | - | - | -2.22 | 0.00 | - | - | - | - | - | - |
| 103 | Taurine and hypotaurine metabolism | - | - | - | - | - | - | - | - | 1.65 | 0.01 | - | - | 1.66 | 0.02 | - | - | - | - | - | - |
| 104 | Bladder cancer | - | - | - | - | - | - | - | - | - | - | -1.68 | 0.05 | -1.64 | 0.04 | -1.53 | 0.04 | -1.73 | 0.02 | - | - |
| 105 | GNRH signaling | - | - | - | - | - | - | - | - | - | - | - | 0. | - | - | - | - | - | - | - | - |

| S. No. | Pathway | Normalized enrichment score | | | | | | | | | | | | | | | | | | | |
|---|---|---|---|---|---|---|---|---|---|---|---|---|---|---|---|---|---|---|---|---|---|
| | | PDR Vs. All control | p-value | PDR Vs. Control (Retina) | p-value | PDR Vs. Control (RPE/C) | p-value | ROP Vs. All control | p-value | nAMD Vs. All control | p-value | nMAD Vs. Control (Retina) | p-value | nAMD Vs. Control (RPE/C) | p-value | nAMD Vs. All dry AMD | p-value | nMAD Vs. Dry AMD (Retina) | p-value | nAMD Vs. Dry AMD (RPE/C) | p-value |
| | pathway | | | | | | | | | | | 1.71 | 03 | | | | | | | | |
| 106 | MAPK signaling pathway | - | - | - | - | - | - | - | - | - | - | -1.74 | 0.02 | - | - | - | - | - | - | - | - |
| 107 | Proximal tubule bicarbonate reclamation | - | - | - | - | - | - | - | - | - | - | 1.54 | 0.03 | 1.79 | 0.01 | 1.92 | 0.00 | 2.01 | 0.00 | - | - |
| 108 | Taste transduction | - | - | - | - | - | - | - | - | - | - | -1.56 | 0.03 | - | - | - | - | 1.49 | 0.04 | - | - |
| 109 | Thyroid cancer | - | - | - | - | - | - | - | - | - | - | -1.66 | 0.02 | -1.88 | 0.00 | - | - | - | - | - | - |
| 110 | Cell adhesion molecules cams | - | - | - | - | - | - | - | - | - | - | - | - | 2.15 | 0.00 | - | - | - | - | - | - |
| 111 | Dilated cardiomyopathy | - | - | - | - | - | - | - | - | - | - | - | - | 1.64 | 0.04 | - | - | -1.5 | 0.04 | - | - |

| S. No. | Pathway | Normalized enrichment score | | | | | | | | | | | | | | | | | | | |
|---|---|---|---|---|---|---|---|---|---|---|---|---|---|---|---|---|---|---|---|---|---|
| | | PDR Vs. All control | p-value | PDR Vs. Control (Retina) | p-value | PDR Vs. Control (RPE/C) | p-value | ROP Vs. All control | p-value | nAMD Vs. All control | p-value | nMAD Vs. Control (Retina) | p-value | nAMD Vs. Control (RPE/C) | p-value | nAMD Vs. All dry AMD | p-value | nMAD Vs. Dry AMD (Retina) | p-value | nAMD Vs. Dry AMD (RPE/C) | p-value |
| | | | | | | | | | | | | | | | | | | 9 | | | |
| 112 | Drug metabolism cytochrome p450 | - | - | - | - | - | - | - | - | - | - | - | - | -1.96 | 0.01 | - | - | - | - | - | - |
| 113 | Fructose and mannose metabolism | - | - | - | - | - | - | - | - | - | - | - | - | -1.70 | 0.01 | - | - | - | - | - | - |
| 114 | Glycosylphosphatidylinositol gpi anchor biosynthesis | - | - | - | - | - | - | - | - | - | - | - | - | -1.74 | 0.01 | - | - | - | - | - | - |
| 115 | Leishmania infection | - | - | - | - | - | - | - | - | - | - | - | - | 2.00 | 0.01 | - | - | - | - | - | - |
| 116 | Metabolism of xenobiotics by cytochrome p450 | - | - | - | - | - | - | - | - | - | - | - | - | -2.19 | 0.00 | - | - | - | - | - | - |

| S. No. | Pathway | Normalized enrichment score | | | | | | | | | | | | | | | | | | | |
|---|---|---|---|---|---|---|---|---|---|---|---|---|---|---|---|---|---|---|---|---|---|
| | | PDR Vs. All control | p-value | PDR Vs. Control (Retina) | p-value | PDR Vs. Control (RPE/C) | p-value | ROP Vs. All control | p-value | nAMD Vs. All control | p-value | nMAD Vs. Control (Retina) | p-value | nAMD Vs. Control (RPE/C) | p-value | nAMD Vs. All dry AMD | p-value | nMAD Vs. Dry AMD (Retina) | p-value | nAMD Vs. Dry AMD (RPE/C) | p-value |
| 117 | Systemic lupus erythematosus | - | - | - | - | - | - | - | - | - | - | - | - | 2.00 | 0.00 | - | - | - | - | - | - |
| 118 | Type I diabetes mellitus | - | - | - | - | - | - | - | - | - | - | - | - | 2.29 | 0.00 | - | - | - | - | - | - |
| 119 | Vascular smooth muscle contraction | - | - | - | - | - | - | - | - | - | - | - | - | 1.88 | 0.00 | - | - | - | - | - | - |
| 120 | RIG I like receptor signaling pathway | - | - | - | - | - | - | - | - | - | - | - | - | - | - | - | - | - | - | - | - |
| 121 | Hypertrophic cardiomyopathy hcm | - | - | - | - | - | - | - | - | - | - | - | - | - | - | - | - | -1.75 | 0.02 | - | - |
| 122 | Aldosterone regulated sodium reabsorption | - | - | - | - | - | - | - | - | - | - | - | - | - | - | - | - | 1.62 | 0.02 | - | - |

| S. No. | Pathway | Normalized enrichment score | | | | | | | | | | | | | | | | | | | |
|---|---|---|---|---|---|---|---|---|---|---|---|---|---|---|---|---|---|---|---|---|---|
| | | PDR Vs. All control | p-value | PDR Vs. Control (Retina) | p-value | PDR Vs. Control (RPE/C) | p-value | ROP Vs. All control | p-value | nAMD Vs. All control | p-value | nMAD Vs. Control (Retina) | p-value | nAMD Vs. Control (RPE/C) | p-value | nAMD Vs. All dry AMD | p-value | nMAD Vs. Dry AMD (Retina) | p-value | nAMD Vs. Dry AMD (RPE/C) | p-value |
| 123 | Basal cell carcinoma | - | - | - | - | - | - | - | - | - | - | - | - | - | - | - | - | -1.61 | 0.03 | - | - |
| 124 | ECM receptor interaction | - | - | - | - | - | - | - | - | - | - | - | - | - | - | - | - | -2.80 | 0.00 | - | - |
| 125 | TGF beta signaling pathway | - | - | - | - | - | - | - | - | - | - | - | - | - | - | - | - | -2.20 | 0.00 | - | - |
| 126 | Chemokine signaling pathway | - | - | - | - | - | - | - | - | - | - | - | - | - | - | - | - | - | - | -1.35 | 0.00 |

**Table S32. Dysregulated pathways of retinal angiogenesis samples compared to controls using RNA-Seq data**

| S.No. | Pathway | NES | NOM p-val |
|---|---|---|---|
| 1 | BLADDER CANCER | 1.78 | 0.02 |
| 2 | PATHWAYS IN CANCER | 1.82 | 0.01 |
| 3 | LYSOSOME | 1.90 | 0.00 |
| 4 | JAK STAT SIGNALING PATHWAY | 1.96 | 0.00 |
| 5 | FC EPSILON RI SIGNALING PATHWAY | 2.04 | 0.01 |
| 6 | HYPERTROPHIC CARDIOMYOPATHY HCM | 2.05 | 0.00 |
| 7 | B CELL RECEPTOR SIGNALING PATHWAY | 2.09 | 0.00 |
| 8 | REGULATION OF ACTIN CYTOSKELETON | 2.09 | 0.00 |
| 9 | FC GAMMA R MEDIATED PHAGOCYTOSIS | 2.11 | 0.00 |
| 10 | SMALL CELL LUNG CANCER | 2.13 | 0.00 |
| 11 | T CELL RECEPTOR SIGNALING PATHWAY | 2.28 | 0.00 |
| 12 | APOPTOSIS | 2.38 | 0.00 |
| 13 | TIGHT JUNCTION | 2.39 | 0.00 |
| 14 | CYTOSOLIC DNA SENSING PATHWAY | 2.59 | 0.00 |
| 15 | OLFACTORY TRANSDUCTION | 2.62 | 0.00 |
| 16 | PRION DISEASES | 2.66 | 0.00 |
| 17 | NOD LIKE RECEPTOR SIGNALING PATHWAY | 2.71 | 0.00 |
| 18 | TOLL LIKE RECEPTOR SIGNALING PATHWAY | 2.76 | 0.00 |
| 19 | SYSTEMIC LUPUS ERYTHEMATOSUS | 2.80 | 0.00 |
| 20 | ECM RECEPTOR INTERACTION | 3.04 | 0.00 |
| 21 | LEUKOCYTE TRANSENDOTHELIAL MIGRATION | 3.06 | 0.00 |
| 22 | NATURAL KILLER CELL MEDIATED CYTOTOXICITY | 3.12 | 0.00 |
| 23 | INTESTINAL IMMUNE NETWORK FOR IGA PRODUCTION | 3.28 | 0.00 |
| 24 | CHEMOKINE SIGNALING PATHWAY | 3.31 | 0.00 |
| 25 | CYTOKINE CYTOKINE RECEPTOR INTERACTION | 3.35 | 0.00 |
| 26 | ASTHMA | 3.42 | 0.00 |
| 27 | ANTIGEN PROCESSING AND PRESENTATION | 3.45 | 0.00 |
| 28 | COMPLEMENT AND COAGULATION CASCADES | 3.46 | 0.00 |
| 29 | AUTOIMMUNE THYROID DISEASE | 3.49 | 0.00 |
| 30 | FOCAL ADHESION | 3.50 | 0.00 |
| 31 | CELL ADHESION MOLECULES CAMS | 3.59 | 0.00 |
| 32 | VIRAL MYOCARDITIS | 3.72 | 0.00 |
| 33 | HEMATOPOIETIC CELL LINEAGE | 3.77 | 0.00 |
| 34 | ALLOGRAFT REJECTION | 3.82 | 0.00 |
| 35 | TYPE I DIABETES MELLITUS | 3.82 | 0.00 |
| 36 | LEISHMANIA INFECTION | 4.03 | 0.00 |
| 37 | GRAFT VERSUS HOST DISEASE | 4.15 | 0.00 |

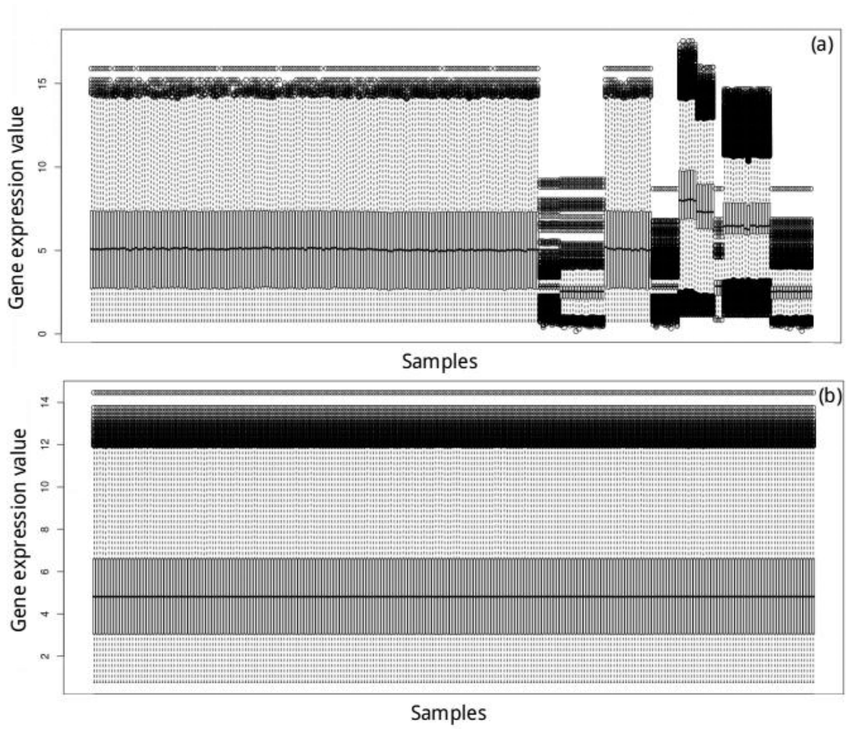


**Figure S1.** Boxplot of expression data (a) before and (b) after quantile normalization

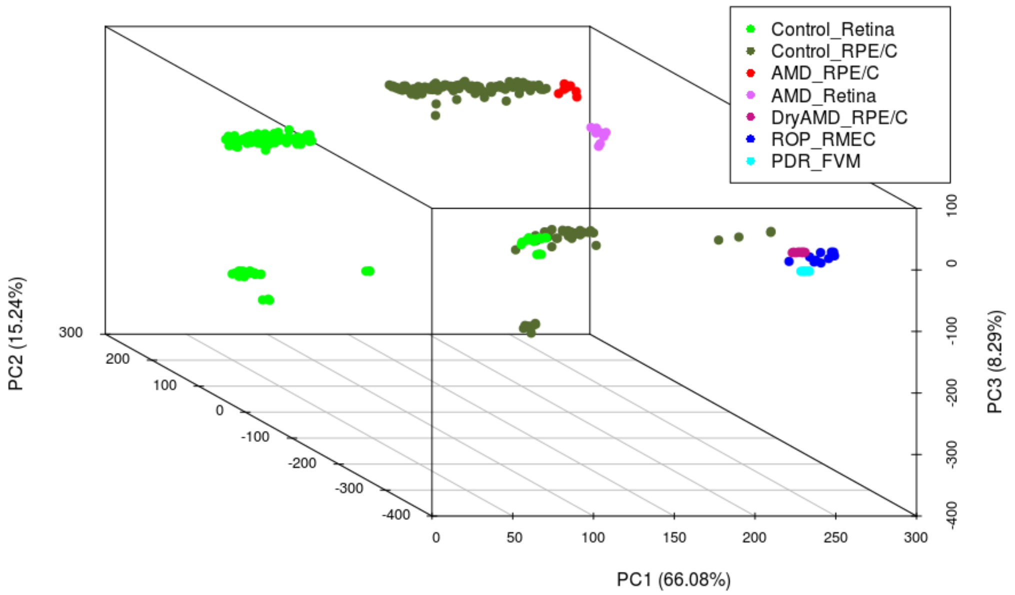


**Figure S2.** PCA plot of samples collected from NCBI-GEO.

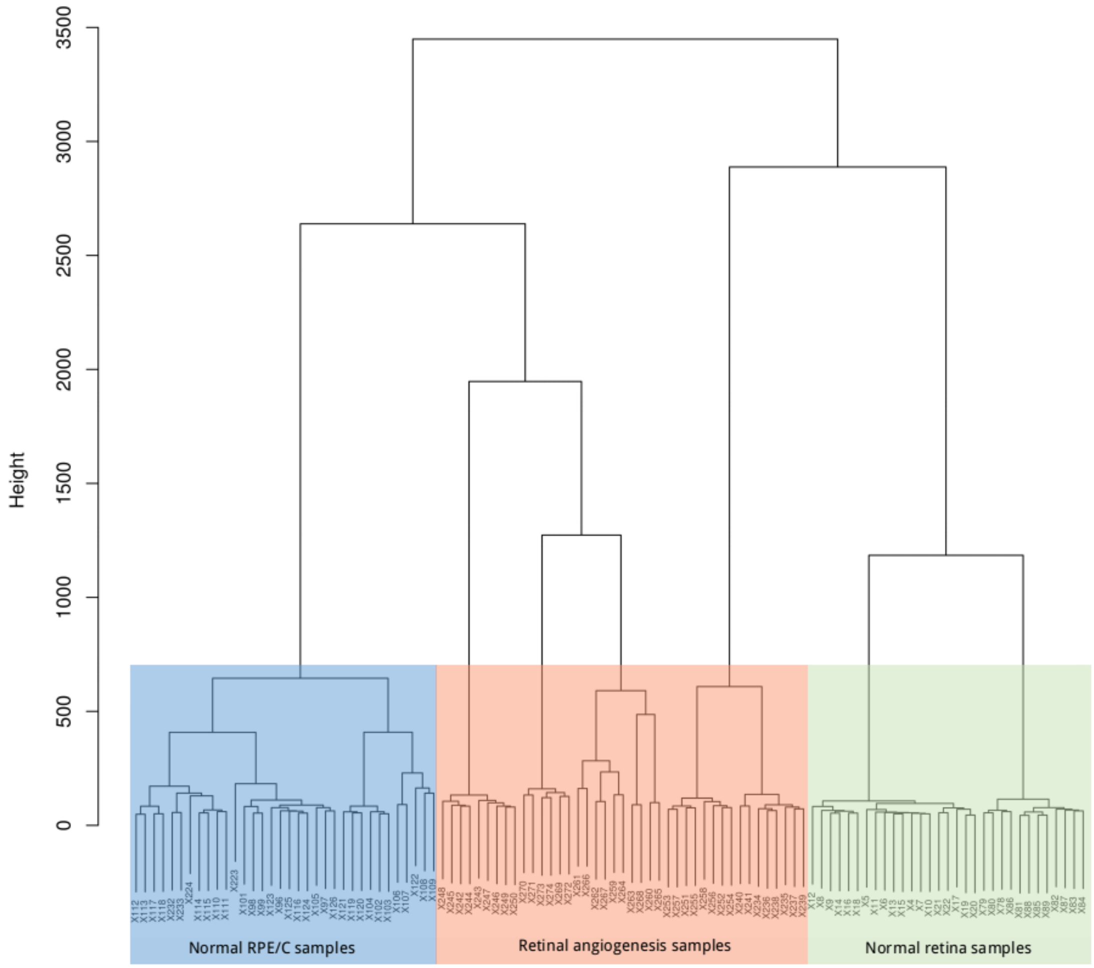


**Figure S3.** Clustering tree after the removal of outlier samples

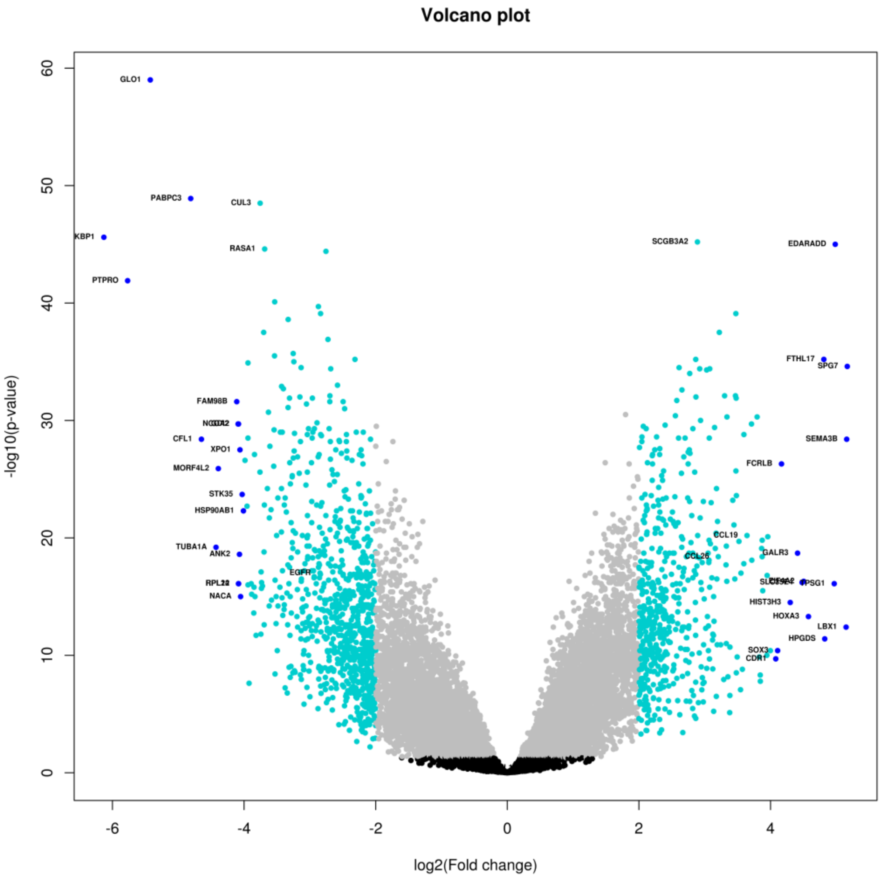


**Figure S4.** Volcano plot of differentially expressed genes in retinal angiogenic disease samples compared to normal control samples (Retina and RPE/C). Black dots are representing genes with no significant p-value and no difference in expression level, gray dots are with significant p-value but no difference in expression level, cyan dots are significantly differentially expressed genes (DEGs), and blue dots are DEGs with high fold change.

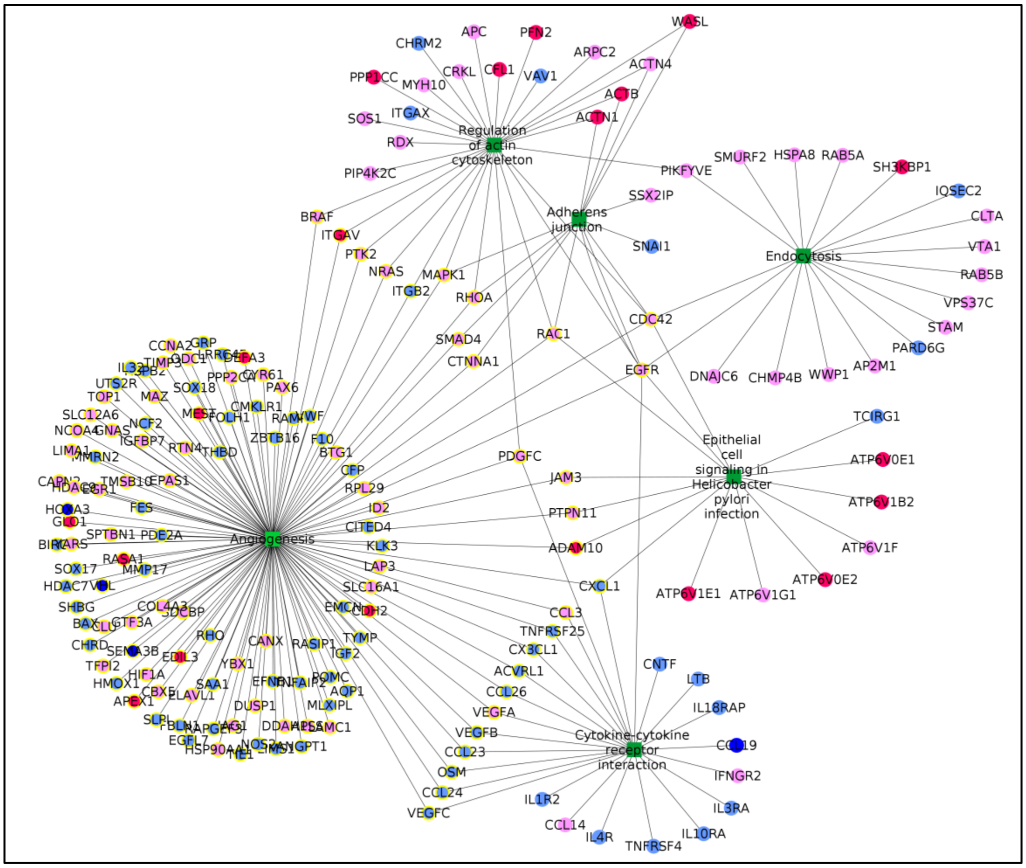


**Figure S5.** Differentially expressed genes in cytokine-cytokine receptor interaction pathway and their association with angiogenesis and other pathways. The green node represents the dysregulated pathways, pink represents DEGs with fold change between -3 and -2, red represents FC ≤ -3, pale blue represents FC between 2 and 3, dark blue represents FC ≥ 3. The round edged square shape of node represents pathways, circle represents DEGs, and yellow circle represents DEGs associated with angiogenesis GO.
